# Local mechanical heterogeneity drives epidermal cell delamination

**DOI:** 10.64898/2026.08.30.747990

**Authors:** Andreas Schoenit, Jérémy O’Byrne, Cécile Daubech, Winfried Schmidt, Lucas Anger, Yuan Shen, Matthias Rübsam, Risai Dubrall, Fanny Wodrascka, Raphaël Voituriez, Benoît Ladoux, Carien M. Niessen, René-Marc Mège

**Affiliations:** Université Paris Cité, CNRS, Institut Jacques Monod, F-75013 Paris, France; Laboratoire Jean Perrin, CNRS, Sorbonne Université, 75005 Paris, France; Max-Planck Zentrum für Physik und Medizin and Max-Planck Institute for the Science of Light, Erlangen, Germany; Department of Physics, Friedrich-Alexander-Universität Erlangen-Nürnberg, Germany; Department Cell Biology of the Skin, Cologne Excellence Cluster on Cellular Stress Responses in Aging Associated Diseases (CECAD), Center for Molecular Medicine Cologne, University Hospital Cologne, University of Cologne, Germany

## Abstract

Delamination within stratified epithelia like the skin epidermis describes the detachment and upward motion of cells originating from the basal layer. Despite its fundamental importance for tissue development, homeostatic regeneration and repair, the mechanisms that drive delamination remain a longstanding open question. Upward motion follows cell shape changes, which are inherently driven by physical forces, but their role is elusive. Here, we investigate delamination in stratifying keratinocytes by combining imaging, force measurements and theoretical modeling. We identify a local change in force balance between differentiating cells and their environment as the key step initiating delamination. Within a homogeneous cell layer with apically polarized contractility, differentiation leads to actomyosin remodeling, redistributing cellular force exertion to the basal side. Such mechanical heterogeneity then results in differentiating cells experiencing and inward basal and outward apical forces that manifest in the formation of a +1 force defect and promote shape changes culminating in upward motion. Simultaneously, delaminating cells actively pull on their underlying neighbors, generating convergent tissue flows which close the basal layer below. Together, we propose a general physical description of delamination initiation, which may act across various multilayered epithelia.

---

Delamination describes the fundamental process of upward cell motion and detachment from the substratum, which underlies the generation and maintenance of stratified epithelia like the epidermis, the outermost layer of the human skin^1–4^. The epidermis protects us from water loss and forms a barrier against various environmental challenges like pathogen infections, UV irradiation, and mechanical deformation^1,2^. This constant exposure to various stresses requires continuous tissue self-renewal, driven by stem cells residing in the basal layer. These stem cells initiate differentiation and move into the upper layers where cells progressively reach terminal differentiation to replace the outermost environment-facing cells that are continuously shed^1–5^.

Early during epidermal development, suprabasal layers are generated by out-of-plane or asymmetric cell divisions^6,7^. In late development and homeostasis after birth, suprabasal layers are maintained by delamination of committed cells, and cell loss in the basal layer is compensated by in-plane cell division^4,8,9^. This differentiation-associated delamination is marked by substantial changes in expression of various adhesion and cytoskeleton proteins like integrins, desmosomal cadherins, or cytokeratins alongside cytoskeletal rearrangements, which are initiated already in the basal layer^9–18^. Such protein expression changes may contribute to cell shape changes and promote delamination^19^. In the mouse ear, long-term intravital imaging of the basal layer showed that cells first initiate differentiation, as demonstrated by the expression of the earliest known differentiation marker Keratin 10 (K10+)^9^, followed by a change in cell shape and finally upward movement^18^. Moreover, shape changes, marked by an increase in apical and decrease in basal area, were shown to be predictive of delamination in the context of tumor initiation^20^. ^9,18^ However, whether and how characteristic shape changes of basal keratinocytes are sufficient to promote upward cell movement, remains unclear^3,19,21^.

More fundamentally, the mechanism underlying this upward movement remains unresolved and has been the subject of longstanding discussions^3–5,21^. Early studies proposed a cell-autonomous, migration-like process^22^. Alternative models suggest that delamination is driven by the local tissue environment, for example through compressive forces^23^, analogous to cell extrusion in simple epithelia^24–26^. Because both cell shape changes within densely packed tissues and cell displacement are inherently mechanical phenomena, understanding the forces involved is essential. Yet, due to the absence of direct measurements of cellular forces or tissue stresses during epidermal delamination, the contribution of mechanics to this process remains unknown.

Here, we investigate the physical principles underlying epidermal cell delamination, focusing on the initiation of shape changes and upward cell movement. To this end, we combined the generation of multilayered epidermal tissues *in vitro* with long-term live imaging, cell-scale force measurements and 3D hydrodynamic modelling. We show that differentiation-associated actomyosin remodeling reshapes the apical and basal distribution of active stresses, giving rise to a mechanical environment that promotes delamination. It consists of traction forces resembling a +1 topological defect, inward-pulling of delaminating cells on their neighbors and convergent tissue flows. Together, these mechanical features drive cell shape changes, initiate upward cell movement and ensure closure of the basal layer. By combining experimental observations with theoretical modeling, we propose a general physical framework for how differentiation generates the mechanical forces drive cell delamination.

## Results

### N/TERT-1 human keratinocytes stratify and form organotypic tissues

We set out to investigate the mechanisms of cell delamination and how an epithelium transits from a monolayer to a multilayer. Human N/TERT-1 basal epidermal keratinocytes^27^ are commonly used to generate organotypic skin equivalents *in vitro*^28^ and thus present a promising model to study delamination. Therefore, we cultivated them in calcium-free growth medium, preventing the formation of cell-cell adhesion, differentiation and stratification^27,28^ (**Figure 1a, ED Figure 1a**). Differentiation was induced at confluence leading to the formation of a first suprabasal cell layer on top of the basal one after 2-3 days (**Figure 1b, ED Figure 1b**). Within five days, multiple suprabasal cell layers were formed (**Figure 1c, ED Figure 1c**). Those layers must have been exclusively generated through delaminations, because in-plane divisions were observed (**ED Figure 1d**). Division-independent stratification suggests homeostatic-like delaminations^3–5^, which are typically preceded by differentiation^18^. Indeed, already one day after differentiation induction, when the tissue was still a monolayer, first basal cells began to express K10 (**Figure 1d**). Moreover, all suprabasal cells observed after two days expressed K10 (**Figure 1e**). When complete stratification was achieved, suprabasal cells showed typical signatures of ongoing differentiation^3–5,10^. Those include a drastic increase in cell size, a change in cell shape and the formation of a lattice-like actin cortex, strong K10 and Dsg1 expression (**ED Figure 1e-g**), resembling *in vivo* and 3D organotypic observations^11,15,29,30^. Thus, our experimental system is biologically relevant and, as it is compatible with live cell imaging, allows to assess cell motion, collective dynamics and cell- and tissue mechanics with high spatiotemporal resolution, which are all inaccessible *in vivo*.

**Figure 1:**
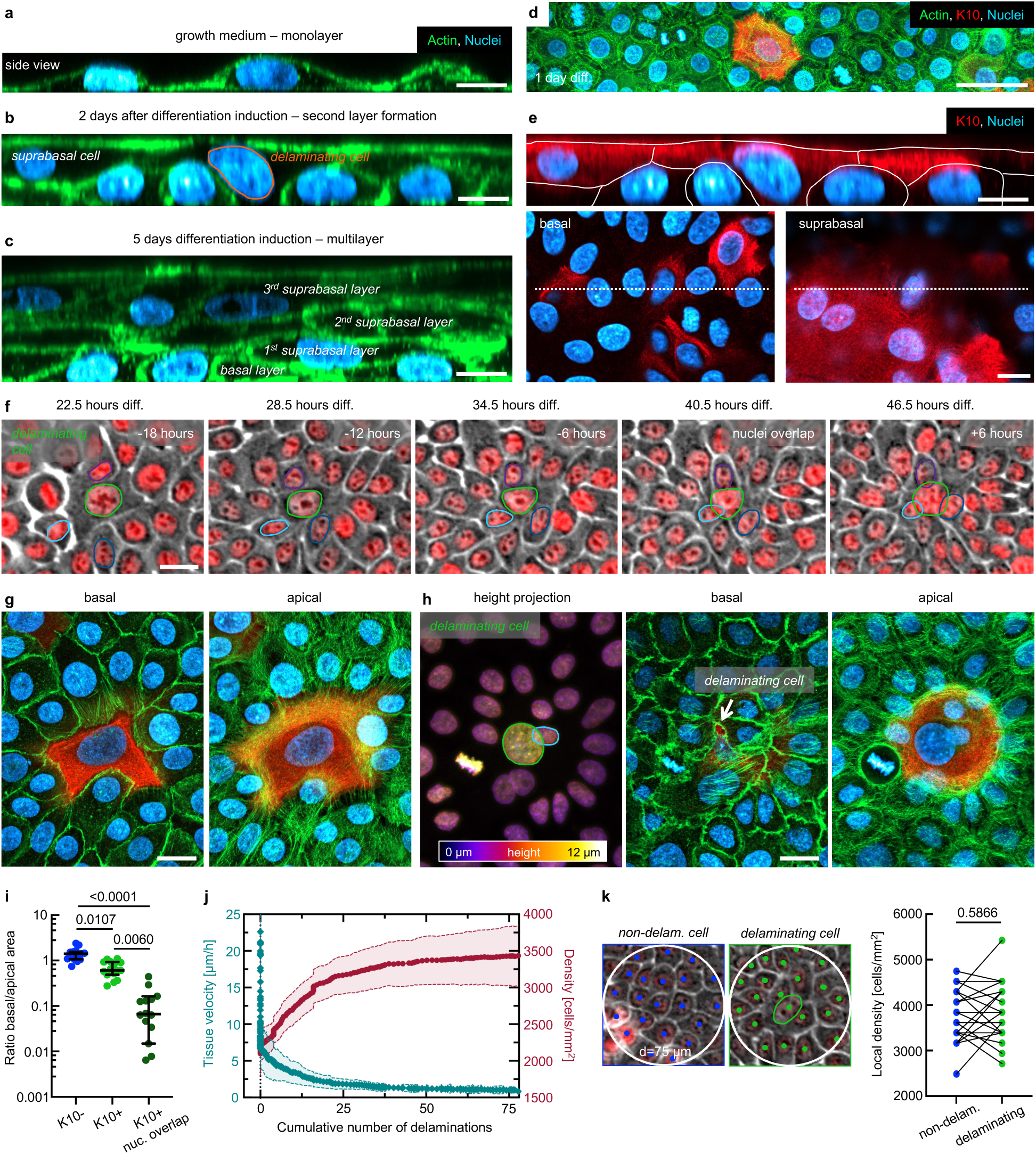
N/TERT-1 basal keratinocytes couple differentiation and delamination. **a** Side view of keratinocytes cultivated in low-calcium growth medium. Nuclei in blue, actin in green. **b** Tissue 2 and **c** 5 days after differentiation induction through a medium switch. Orange outline shows a delaminating cell. **d** XY view 1 day after differentiation induction. K10 in red. **e** Top: Side view of K10 expression after 2 days, corresponding to **b**. White lines show cell borders, based on actin staining in **b**. Bottom: XY view of the basal (left) and the suprabasal plane (right). Dotted line indicates the position of the side view. **f** Phase contrast time sequence of a detected delamination event, which is defined as the first nuclei overlap with one neighbor cell. Timestamps correspond to time after differentiation induction (black), or to time relative to nuclei overlap (t=0 hours, white). **g** Confocal images of a K10+ cell and its surrounding before nuclei overlap. Apical (left) and basal (right) view. **h** Delaminating K10+ cell after nuclei overlap. Left: Height-projection showing the height differences between the delaminating cell (yellow) and basal cells (purple). Middle: Apical view of the delaminating cell and the surrounding tissue. Right: Basal view. Arrow indicates remaining basal adhesive area. **i** Quantification of the ratio between basal and apical area, corresponding to **g** and **h**. Mean +/− SD from N=5 independent experiments and n=15 cells. P-values from multiple ANOVA test. **j** Tissue velocity (cyan, left Y-axis) and cell density (red, right Y-axis) as a function of the cumulative number of tracked delaminations. Mean +/− SD from N=3 independent experiments and n=78 events. **k** Phase contrast (left) and quantification (right) of the local cell density within a 75µm circle at t=0 hours within the same frame. P-value from paired t-test from N=3 independent experiments and n=20 events. Scale bars 10 µm (**a**, **b**, **c**, **e**); 20 µm (**f**, **g**, **h**); 50 µm (**d**).

### Cell delamination is coupled to differentiation

We used long-term live cell imaging to continuously track delaminating cells. Those became visible the second day after differentiation induction and were identified by the partial overlap of two nuclei, which gradually increased during the observation period (**Figure 1f**, *Supplementary Movie 1*). Such cells did not divide over 48 hours, indicating that they had differentiated and exited the cell cycle^3^. *In vivo,* the complete delamination process takes several days and involves progressive basal shrinking and simultaneous apical expansion^18^. To investigate the state of delamination when nuclei partially overlapped, we analyzed fixed tissues and assessed the ratio between basal and apical areas (**Figure 1g-i**). Non-differentiated cells showed on average a ratio close to one. (**Figure 1i**). K10+ cells already showed a larger apical area before nuclear overlap. The basal-to-apical ratio was strongly decreased when nuclei partially overlapped, as only a small basal cell-substrate interface remained (**Figure 1i**). These ratios were in good agreement with *in vivo* measurements reported by Cockburn *et al.*^18^ at the final stage of delamination (**ED Figure 2a**). Thus, the nuclei overlap will serve as a 2D proxy for late delamination and as the reference timepoint (t = 0 hours) to align delamination events for all further analysis. For every tracked delaminating cell, we tracked as an internal control random non-delaminating cells that divided at least once during the observation period (**ED Figure 2b**). Those were assigned the same reference timepoints as the delaminating cells (**ED Figure 2b**) to correct for time- and calcium-dependent global tissue changes.

### Delaminations follow a tissue jamming transition and density increase

We next investigated how the onset of delamination relates to the evolving tissue state. By quantifying global cell velocities, we identified a jamming transition within the first day upon differentiation induction (**ED Figure 2c**, *Supplementary Movie 2*). The first delamination events occurred subsequently (**ED Figure 2c**), consistent with previous reports^23^. This jamming transition may result from the progressive establishment of cadherin-mediated cell-cell adhesion promoted by the high Ca^2+^-concentration that is present in the differentiation medium. Cell-cell contacts initially formed between pairs of cells or within small clusters, and progressively expanded to eventually span the entire system, resembling an adhesion percolation process^31^ (**ED Figure 2d**). Interestingly, this process was accompanied by a strong increase in global tissue stiffness, indicating a rigidity transition (**ED Figure 2e**).

We next assessed global cell density, which increased initially and then plateaued despite continuous cell divisions, as these were counter-balanced by delaminating cells that left the basal layer (**ED Figure 2f**), and most delaminations occurred when the density had plateaued, correlating with a jammed tissue state (**Figure 1j**). Such conditions are seen in late development^8^ and in a homeostatic epidermis^32^, where however delamination may be triggered by local rather than tissue-wide changes. A transient increase in local cell density caused by neighboring cell divisions may promote delamination^23^. We therefore compared the number of neighbor divisions (**ED Figure 2g**) and local cell-density fluctuations (**Figure 1k**) near non-delaminating and delaminating cells. Neither parameter differed significantly between the two groups, indicating that local crowding alone is insufficient to explain delamination and pointing instead to additional physical determinants.

### Delaminating cells increase in size, correlating with a tissue organization defect

A longstanding body of work has linked differentiation to an increase in cell size^33–35^. Consistent with this, K10+ N/TERT-1 keratinocytes already displayed an enlarged area one day after differentiation induction (**ED Figure 3a**). We therefore examined the morphology of the earliest delaminating cells and found that, as early as 24 hours before nuclei overlap, they exhibited significantly larger cell and nuclei areas than non-delaminating cells (**Figure 2a,b, ED Figure 3b**). This emerging cell size heterogeneity was accompanied by a marked reorganization of the local epithelial topology. Whereas non-delaminating cells maintained six neighbors on average, cells destined to delaminate progressively gained neighbors, reaching an average of ten at the time of nuclei overlap (**Figure 2c**). Thus, excessive cell and nuclear growth is coupled to a pronounced tissue organization defect, providing an early and robust structural signature of impending delamination

**Figure 2:**
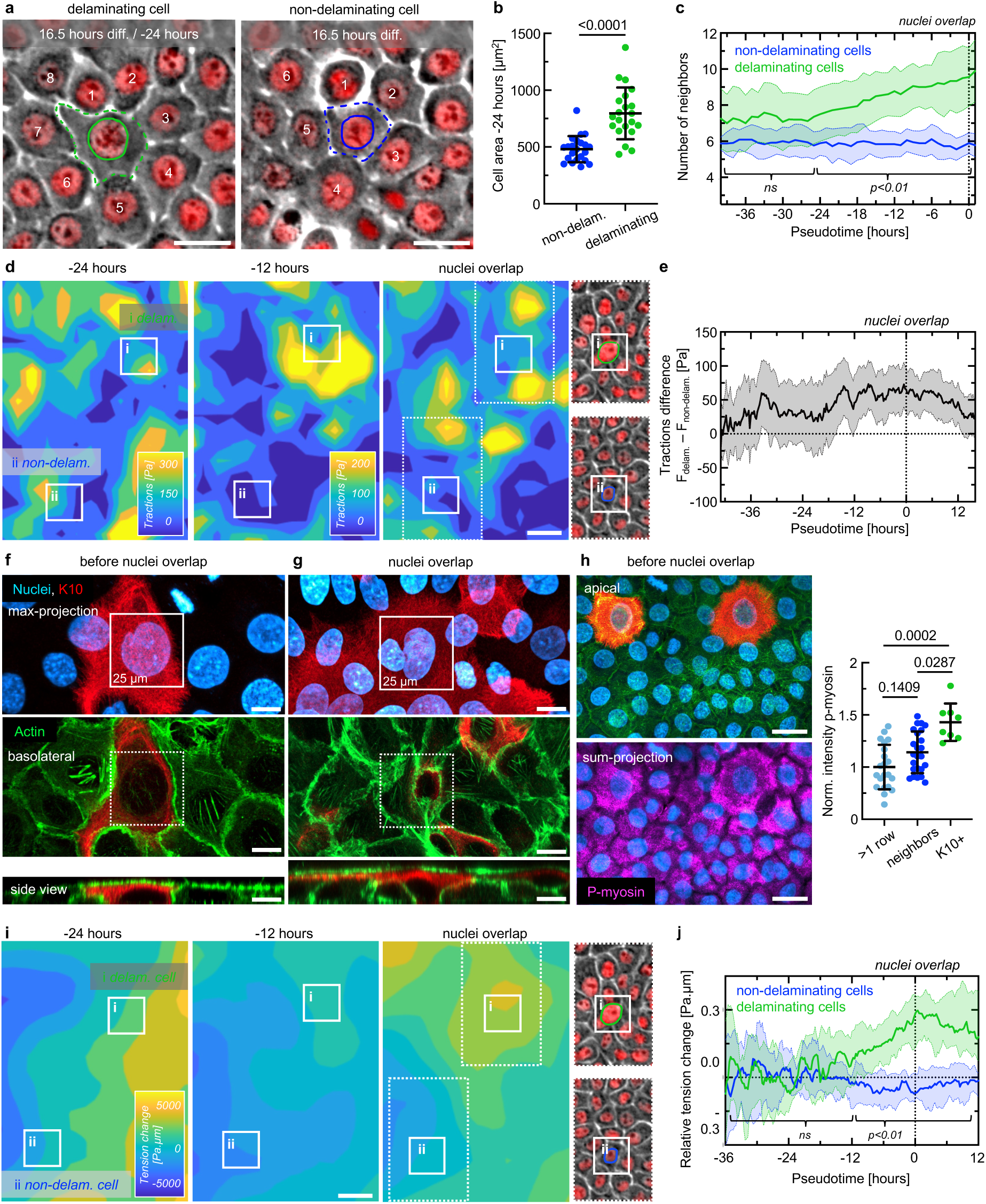
Increased local traction force generation and tissue tension around delaminating cells. **a** Phase contrast images of non-delaminating and delaminating cells 24 hours before nuclei overlap. Manual annotation of the cell outlines and the neighbors. **b** Quantification of the cell area 24 hours before nuclei overlap. Means +/− SD. P-value from t-test. N=2 independent experiments, n=22 cells. **c** Number of neighbors through time relative to nuclei overlap at t=0. Median +/− 95% CI of n=18 individually tracked cells from N=3 independent experiments. **d** Example traction force fields relative to nuclei overlap. White windows show local areas of measurement for (i) delaminating and (ii) non-delaminating cells. White dotted lines show phase contrast image at nuclei overlap. Same scale at −12 hours and nuclei overlap. **e** Quantification of traction force difference (F_delaminating_ – F_non-delaminating_) relative to nuclei overlap. Median +/− 95% CI of n=67 individually tracked cells from N=3 independent experiments. **f** Confocal images of a K10+ (red) cell long before the nuclei overlap and **g** close to it. *Top*: maximum-projection nuclei (blue) and K10. *Middle*: basolateral view of actin cytoskeleton (green). *Bottom*: side views. White and dotted boxes show the size of traction force quantification related to **d**, **e**. **h** Confocal images 1 day after differentiation induction. *Top*: apical view. *Bottom*: Sum-projection over the whole cell volume showing phospho-myosin (magenta). Right: quantification of the phospo-myosin intensity in K10+ cells, their neighbors and cells more than one row away from K10+ cells. Normalization to the average of the cells more than one row away. Means +/− SD. Each datapoint shows one cell from N=2 independent experiments. **i** Stress maps indicating relative tension changes at different timepoints. Boxes show areas of (i) delaminating and (ii) non-delaminating cells. Dotted lines show corresponding phase contrast and nuclei (red) image. **j** Quantification of the relative tension change in positions as indicated in **i** leading to the nuclei overlap. Median +/− 95% CI of n=67 individually tracked cells from N=3 independent experiments. p-value from continuous t-test; brackets indicate timeframe where p<0.01 or not significant (ns). Scale bars 10 µm (**f**, **g**); 25 µm (**a**, **d**, **h**, **i**).

### Local traction forces increase before nuclei overlap

Heterogeneous tissues often show cell-type specific differences in force generation, as observed in the context of cell competition^25^, within intestinal organoids^36^ and *in vivo* during skin placode formation^37^. A key advantage of our *in vitro* model is its compatibility with traction force microscopy^38^ to track how changes in the mechanical environment relate to alterations in cellular force generation. We investigated traction forces globally and on the scale of single cells (within a 25 µm window, **Figure 2d**). Because the tissue jamming transition is accompanied by an overall global traction force decrease (**ED Figure 3c,d**), likely due to increasing cell-cell^39^ or changing cell-substrate adhesion^40^, we report the difference in force magnitude around delaminating and non-delaminating cells (**Figure 2e**). Traction forces were significantly elevated around cells destined to delaminate as early as 18 hours before nuclei overlap. This increase persisted for more than a day before decreasing again several hours after nuclei overlap (**ED Figure 3d**).

Delamination is accompanied by strong cell shape changes. Because the analysis window size approximates typical basal areas of K10+ cells (**Figure 2f**), measurements long before nuclei overlap thus captured forces likely generated mainly by the delaminating cells, and suggests that force production increases as part of the differentiation program. As nuclei overlap approached, the measured traction increasingly reflected forces generated by neighbor cells as they expanded beneath the delaminating cell (**Figure 2g**). Radially averaging at the time of nuclei overlap revealed a pronounced force maximum approximatively 25 µm away from the delaminating nucleus (**ED Figure 3e**). Beyond this distance, traction rapidly declined, indicating that delamination is driven by a spatially confined mechanical unit composed of the delaminating cell and its immediate neighbors, with little contribution from more distant cell rows.

### Local cell contractility and tissue tension increase during delamination

Because elevated traction forces are generally associated with increased cellular contractility^25^, we next compared total phospho-myosin levels between basal K10+ and K10-cells one day after differentiation induction. Confirming TFM measurements, K10+ cells showed a significant increase in total phospho-myosin staining compared to distant cells (**Figure 2h**), in agreement with *in vivo* reports of elevated contractility in suprabasal cells^23,29,41^. Immediate neighbors of K10+ cells also showed a trend towards increased phospho-myosin levels, suggesting that the mechanical response extends beyond the differentiating cell itself, in line with traction forces (**Figure 2h**). This mechanically distinct domain is spatially restricted to the future delaminating cell and its first row of neighbors, indicating that differentiation locally generates mechanical heterogeneity.

Heterogeneity in traction force magnitude is associated with local differences in intercellular stresses^25^. Thus, we next asked whether tissue stresses are dynamically remodeled during delamination. Laser ablation first established that the epithelium was globally under tension (**ED Figure 4a-c**, *Supplementary Movie 3*), in line with previous reports^15^. To monitor relative tension fluctuations over time, we used Bayesian Inversion Stress Microscopy (BISM)^42,43^. Whereas tension remained constant around non-delaminating cells, cells destined to delaminate exhibited a gradual tension increase starting around 12 hours before nuclei overlap (**Figure 2i,j**, *Supplementary Movie 4*). Excess tension gradually declined after overlap but remained elevated for more than 12 additional hours. This shows that delamination is accompanied by a sustained, highly localized tensile state lasting more than one day. Together, our results establish a previously unrecognized link between differentiation-associate actomyosin activation and an increase in traction forces and tensile stresses that preceded delamination. These mechanical features may actively create the physical conditions required to initiate and drive delamination.

### Delamination is accompanied by the formation of a +1 traction force defect

To determine how the elevated local traction forces organize over time, we investigated their orientation (**Figure 3a**, *Supplementary Movie 5*). We quantified force convergence - the negative divergence - relative to the center of tracked cells. Initially, traction forces showed no preferred orientation around either delaminating or non-delaminating cells.

**Figure 3:**
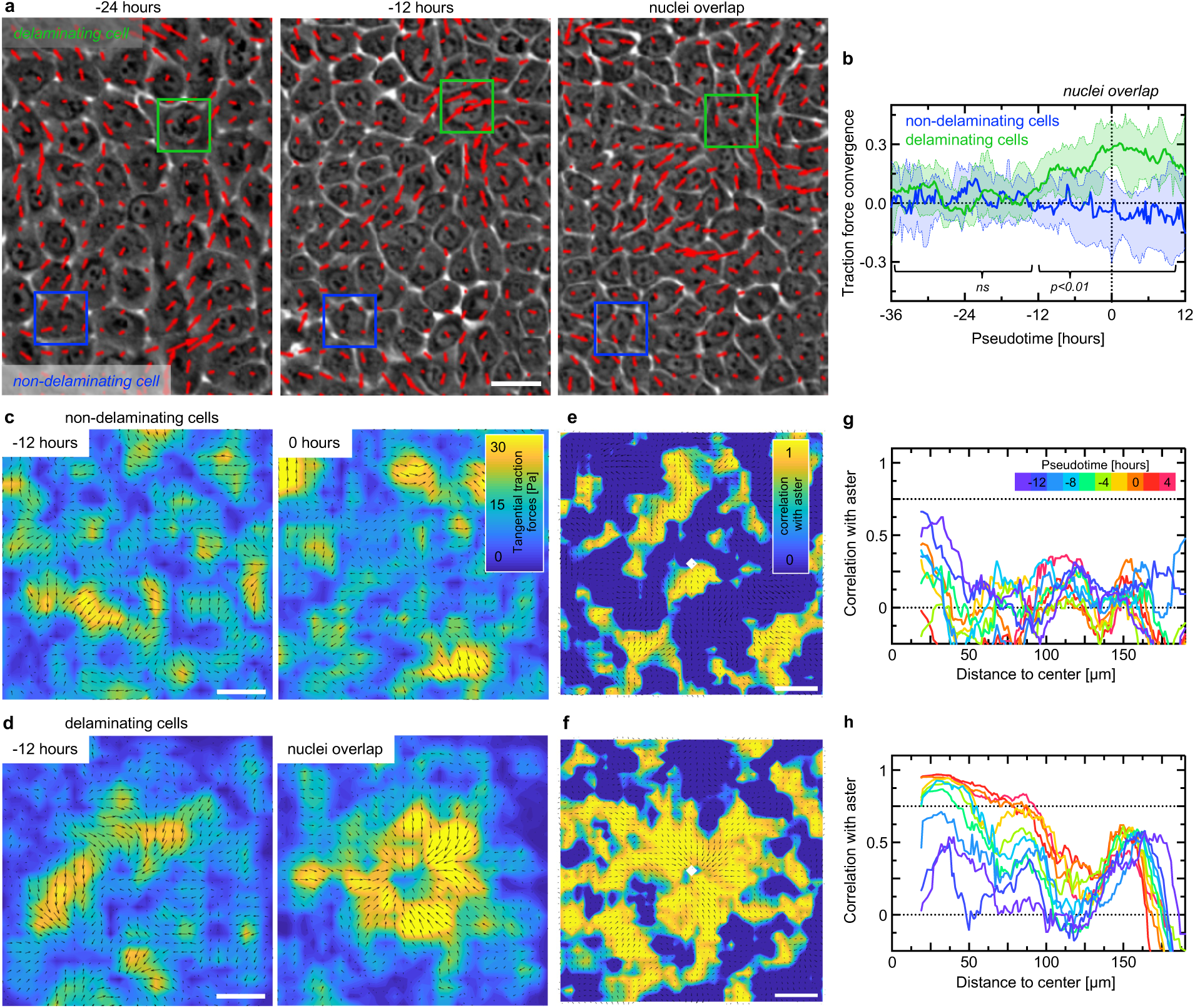
Emergence of a +1 traction force defect. **a** Phase contrast images overlayed with traction vectors normalized to the highest vector in the field at different timepoints. Boxes indicate areas of local orientation quantification. **b** Temporal evolution of the traction force convergence (inward orientation) relative to nuclei overlap. Median +/− 95% CI of n=67 individually tracked cells from N=3 independent experiments. **c** Averaged traction force field around n=67 non-delaminating and **d** delaminating cells 12 hours before and at the nuclei overlap. **e** Correlation map of experimental traction force field and aster pattern for non-delaminating and **f** delaminating cells at nuclei overlap. **g** Radially averaged correlation as a function of the distance to the center around non-delaminating and **h** delaminating cells. The time is indicated by progressively warm colors. p-value from continuous t-test; brackets indicate timeframe where p<0.01 or not significant (ns). Scale bars 25 µm (**a**, **c**); 50 µm (**e**).

Approximatively 12 hours before nuclei overlap, however, forces surrounding future delaminating cells began to orient inward and progressively converged towards the cell center. This convergent organization strengthened over time and persisted until several hours after nuclei overlap (**Figure 3b**). Spatially averaged force fields further revealed a striking difference between the two cell populations. Around non-delaminating cells, traction forces remained disordered at all timepoints, with no reproducible spatial organization (**Figure 3c**, *Supplementary Movie 6*). By contrast, forces around delaminating cells self-organized into an inward-pointing pattern, which became particularly pronounced at nuclei overlap (**Figure 3d**, *Supplementary Movie 7*). Within the framework of active matter^44^, this emergent pattern corresponds to an aster-like +1 topological defect. Our findings therefore identify delamination as a process associated not only with elevated force magnitude, but with the emergence of a highly ordered topological force pattern that may act as a local organizer of epithelial remodeling^45^.

To further quantify the emergence and spatial extend of this defect, we generated correlation maps representing the similarity between a simulated aster and the experimental force pattern (**Fig. 3e,f, ED Figure 3f**) and then radially averaged the correlation through time. Due to random fluctuations in force direction, non-delaminating cells showed a correlation that did not exceed 0.65 (**Figure 3g**). A similar correlation was observed for delaminating cells at early timepoints. Progressing towards nuclei overlap, however, the correlation approached 1 in a wide radius around the delaminating cell (**Figure 3h**). The spatial extent of this pattern also increased over time. Correlation values above 0.75 extended over a radial distance exceeding 50 µm near nuclei overlap, corresponding to a defect diameter greater than 100 µm (**Figure 3h**). This was nearly twice the size of the typical apical diameter of a delaminating cell after two days of differentiation (**ED Figure 4d,e**) and approximately matched the combined dimension of the delaminating cell and its first row of neighbors. These findings demonstrate that the topological force defect consists of a multicellular structure rather than a cell-autonomous feature and further highlight the immediate neighboring cells as integral mechanical contributors to the delamination process.

### Actomyosin remodeling is associated with differentiation

Having uncovered a traction force defect as a defining mechanical feature of delamination enabled us to revisit the question of the physical mechanisms that drive cell shape changes and promote delamination^18,20^. One possibility is that neighboring cells protrude beneath the delaminating cell, thereby reducing its basal area. This mechanism is unlikely, however, because this would typically generate traction force patterns opposite from what is observed here, with forces oriented in opposite direction of protrusion^46^.

Differentiation-induced actin remodeling was shown to be crucial for delamination^14,29^. We therefore asked whether delaminating cells expand apically over their neighbors through lamellipodia-like protrusions. Long-term live-cell imaging revealed no evidence of such protrusions. Instead, it uncovered a pronounced remodeling of the actin cortex (**ED Figure 5a**, *Supplementary Movie 8*).

To determine how actin remodeling could drive apical expansion, we assessed more closely subcellular changes in actomyosin architecture, and abundance in the apical and basal planes. As observed in other basal keratinocytes^30,47^, undifferentiated N/TERT-1 cells formed in the apical plane a highly ordered interconnected actin network (**Figure 4a**) upon engaging in cell-cell adhesion. Phospho-myosin clustered apically at the nodes where stress fibers converged on top of the nuclei (**Figure 4b**). By contrast, K10+ cells displayed a profoundly reorganized actin cytoskeleton, characterized by large orthoradial actin arcs enriched in phospho-myosin (**Figure 4c,d**). Actin levels per unit area were slightly reduced in K10+ cells, while phospho-myosin levels were increased (**Figure 2**, **ED Figure 5b**). Moreover, actomyosin was redistributed towards the cell periphery, where the arcs were most prominent, in marked contrast to the central actomyosin clusters observed in K10-cells (**Figure 4e**).

**Figure 4:**
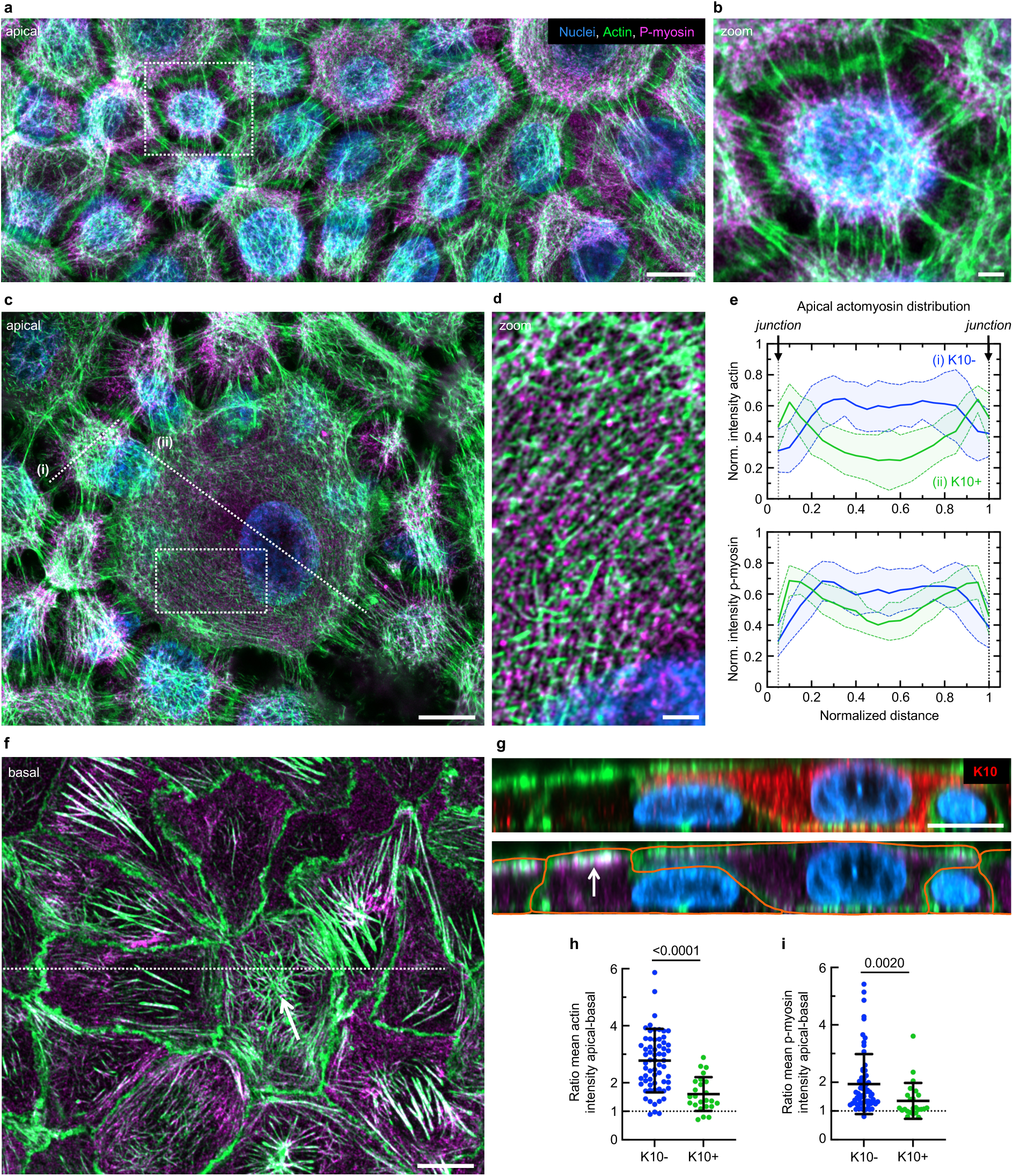
Differentiation-induced actomyosin redistribution. **a** Representative confocal image of the apical actomyosin cytoskeleton in undifferentiated, non-delaminating cells. Actin in green, phospho-myosin in magenta, nuclei in blue. **b** Zoom-in at position indicated in **a**. **c** Apical view of the actomyosin distribution in a delaminating cell and its neighborhood. **d** Magnified and rotated view of the apical actomyosin network in the delaminating cell at the position indicated in **c**. **e** Horizontal distribution of actin (top) and phospho-myosin (bottom) across the apical planes of (i) K10- and (ii) K10+, as indicated by dotted lines in **c**. Mean +/− SD from n=25 cells across N=3 independent experiments. **f** Basal view of actomyosin corresponding to **c**. White arrow indicates radial actin accumulation in the basal center of the delaminating cell, below its nucleus. **g** Side views showing the apical expansion of a differentiating cell on its neighbors. Top: K10 (red). Bottom: Lateral actomyosin distribution. Cell outlines drawn in orange. White arrow shows pronounced apical actomyosin accumulation in neighbor cells. **h** Ratio of the mean actin and **i** phospho-myosin intensity per unit area comparing the apical (**c**) and basal (**f**) planes. Each datapoint shows one cell from N=3 independent experiments. Scale bars 2 µm (**b**, **d**); 10 µm(**a**, **c**, **f**, **g**).

In the basal plane, actomyosin levels per unit area were not only elevated in K10+ cells (**ED Figure 5c**) but its organization was also extensively remodeled. In K10+ cells, actin fibers accumulated isotropically beneath the nucleus, whereas neighbor cells formed stress fibers preferentially oriented radially toward the center of the delaminating cell (**Figure 4f**, **ED Figure 5d**). The subcellular phospho-myosin distribution remained comparatively uniform in both populations (**ED Figure 5e**). Importantly, quantifying the vertical distribution revealed a strongly decreased apical-basal polarization of actin and myosin within K10+ cells (**Figure 4g-i**). Together, these findings show that differentiation not only increases overall phospho-myosin levels but also triggers a comprehensive reorganization of the force-generating cytoskeleton. Actomyosin shifts from a centrally focused, apically polarized network to a more peripheral apical architecture accompanied by an increase in an isotropically organized basal network. This vertical and radial redistribution provides a structural basis for the observed changes in force transmission and for alterations in cell shape that promote delamination.

### Theoretical model of differentiation-induced shape changes

Based on our observations that differentiated cells display a significant reorganization of their actomyosin cytoskeleton both at their basal and apical sides, we hypothesized that the local mechanical heterogeneity induced by the differentiating cell in the monolayer is sufficient to trigger delamination. To substantiate our hypothesis, we developed a minimal, active hydrodynamic model of a cell monolayer hosting a delamination event. The key ingredient of the model is that, because of its differentiated state, the cell undergoing delamination imposes specific mechanical boundary conditions to its neighbors, which are assumed to remain in a non-differentiated homeostatic state. More explicitly, we consider a single differentiating cell, within a uniform monolayer of otherwise undifferentiated cells. We adopt a phenomenological, hydrodynamic description of the monolayer, modelled as a uniform, isotropic, incompressible active fluid layer of viscosity η^48–50^. This is justified by the slow timescales of delamination^18^. The delaminating cell is assumed to remain axisymmetric over the full process and is defined by its boundary r_c_(z) in standard (r,θ,z) cylindrical coordinates. We assume that the delaminating cell starts with a columnar shape, r_c_(t = 0,z) = R, within a monolayer at rest with uniform thickness h, where we take h = R for simplicity. Our key assumption is that the differentiation process imposes distinct rheological and mechanical properties for the delaminating cell, and particularly a different, z-dependent active stress **τ**, which accounts for the observed vertical actomyosin redistribution. For simplicity, we assume that other parameters remain unchanged. Those include viscosity and friction properties with the substrate, which define the friction length l_f_ (see below). With these definitions, the merit of the model is to minimally account for the differentiation process by imposing the active stress difference Δ**τ**(z) at the boundary between the delaminating cell and its neighbors (**Figure 5a**).

**Figure 5:**
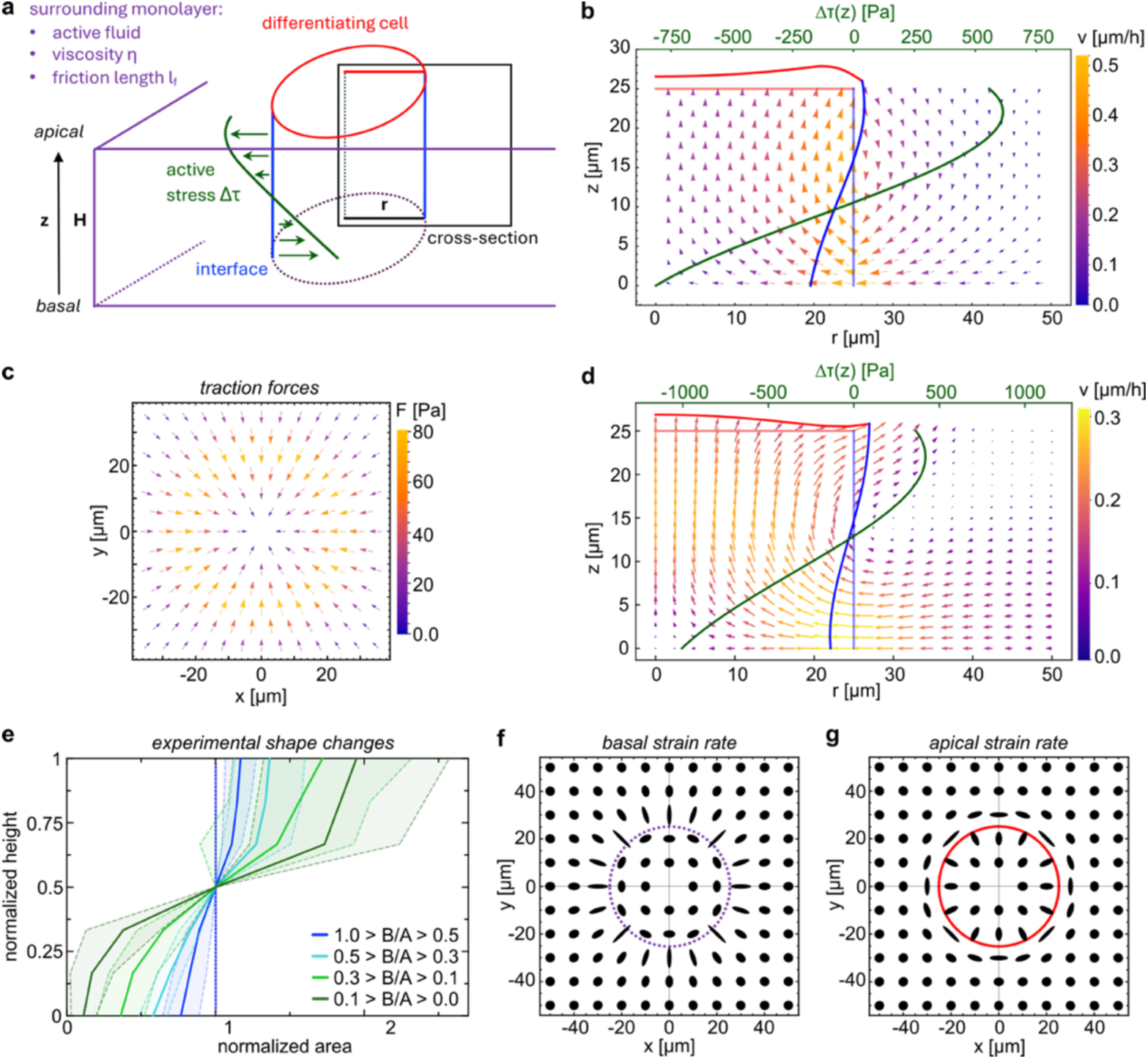
A theoretical model shows how differences in active stress promote cell shape changes. **a** Schematic of the hydrodynamic model. The tissue of height z is modeled as an active fluid with fixed viscosity and friction with the substrate. It contains a cylindrical differentiating cell with radius r. Differentiation is minimally modeled by a differential active stress amplitude Δτ (green line) along the vertical axis of the interface with its surrounding (blue line). As it remains axisymmetric, deformations are investigated within a cross-section (black window). **b** Resulting flow field. The green line is the jump of the active stress amplitude Δτ(z) between the delaminating cell and the rest of the tissue. The displayed vector field is the velocity field resulting from this stress imbalance. In turn, this flow advects the delaminating cell’s boundary (pale blue and red lines at t=0; darker blue and red at time t∼10 hours). **c** The corresponding traction force field exerted on the substrate. **d** Numerical simulations of the flow field and resulting cell shape changes at time t=0 (pale blue and red lines) and t∼10 hours (dark blue and red lines). **e** Experimentally measured cell shape changes of differentiated K10+ cells. The basal-to-apical area ratio (B/A) serves as a proxy of delamination progression, where the basal area is approaching zero. Normalization to maximal value. All curves show means +/− SD. **f** Strain rate projected on the apical and **g** basal xy planes. Circles indicate dimensions of the delaminating cell.

The problem can then be formulated explicitly as a heterogeneous active Stokes fluid with a dynamic boundary, on which, according to Δ**τ**(z), the delaminating cell is pulling stronger on the basal side, while the surrounding tissue pulls stronger on the apical side (see *Supplementary Information*). This allows to obtain the flow field **v**(r,z) (**Figure 5b**), which grants access to several observables that can be measured experimentally, including the traction force pattern (**Figure 5c**). We combined this analytical model with a numerical approach (**Figure 5d**) to access different possible boundary conditions and fitted two independent parameters: first, the friction length l_f_ ≈ 20.9 µm, based on traction force measurements (**ED Figure 6a**), which allowed obtaining the fluid viscosity η_eff_ ≈ 4 kPa/h (**ED Figure 6b**). Second, the typical delamination speed v_typ_ ≈ 0.5 µm/h, based on cell size and the timescale of mechanical changes reported above, which agrees with *in vivo* observations^18^. The joint analytical and numerical model recapitulates the early-time dynamics of the delamination process, including the 3D shape changes of the delaminating cell parametrized by r_c_(z,t) and h(r,t) (**Figure 5b,d**). These predictions are consistent with experimental observations during the early stages of delamination (**Figure 5e**).

Using the numerical model (**Figure 5e**), we individually shifted l_f_ and found that a more than 2-fold decrease or increase (10 or 50 µm, respectively) changed the flow pattern quantitatively and thus only the magnitude of typical shape changes (**ED Figure 6c,d**). In contrast, changing the mean, z-averaged active stress difference Δ**τ** results in qualitative flow pattern changes and consequently different cell shapes. A negative Δ**τ** indicates that the differentiating cell pulls overall stronger than the surrounding tissue, resulting in an increased cell height and a more columnar shape (**ED Figure 6e**). A positive Δ**τ** indicates stronger pulling of the surrounding tissue in the apical plane, which results in vertical stretching and a decreased height of the differentiating cell (**ED Figure 6f**). Thus, within the explored parameter space, the z-averaged active stress difference determines the resulting qualitative cell deformations. It suggests that a disturbed force balance could hinder delamination by generating shapes which may prevent upward motion.

Investigating the predicted strain rates in the basal and apical planes revealed strikingly different patterns at the heterotypic boundary: basally, the neighbors deform orthogonal to the delaminating cell (**Figure 5f**), but apically, their deformations show an orthoradial orientation (**Figure 5g**). Those strain rates resemble the experimentally observed shapes and particularly the organization of the actin cytoskeleton (**Figure 4**), thereby linking the measured cytoskeletal architecture to the underlying tissue-scale mechanics. Importantly, the model extended our experimental, 2D in-plane tension measurements by resolving how stresses vary along the apicobasal axis. The imposed active stress difference Δ**τ**(z) leads to a z-dependent switch from basally compressive to apically tensile stresses at the interface (**ED Figure 6g**). Thus, although our experimental data revealed an increase in in-plane tensile stress, the model predicts that the tensile state coexists with a spatially confined basal compression. This three-dimensional stress architecture provides a mechanical explanation for the simultaneous basal constriction, apical expansion, and upward displacement that characterize delamination. Finally, we show that our analysis can also consistently predict the emergence of nematic order: assuming that the monolayer is in the isotropic phase at rest before differentiation starts, we find that the classical flow alignment coupling^48–50^ then leads to the emergence of local nematic order characterized by the orientational order parameter (Q-tensor) around the delaminating cell, with the symmetry of a +1 topological defect (see *Supplementary Information*). Altogether, our analysis shows that the mechanical heterogeneity initiated by the differentiating cell, minimally described by a z-dependent difference in active stress, is sufficient to explain delamination-initiating shape changes, will impose a local +1 topological defect structure, and predicts convergent flows of the surrounding tissue.

### Delamination generates a convergent tissue flow

A central prediction of our theoretical model is the emergence of convergent tissue flows due to the differentiation-driven vertical redistribution of active stresses. To test this prediction, we tracked cells around delaminating cells. Indeed, we observed that neighbor nuclei slowly moved towards the delaminating cell over the course of several hours before the nuclei overlap (**Figure 6a**, *Supplementary Movie 9*). We quantified these local tissue movements by measuring flow orientation within a 50-µm window centered on delaminating cells using particle image velocimetry (PIV) (**ED Figure 7a**). At early timepoints, flows showed no preferred orientation. Approximately 12 hours before nuclei overlap, however, they became significantly convergent and remained so for more than one day. No comparable convergence was detected around non-delaminating cells (**Figure 6b**). Such flows can arise from collective cell migration^46^ and have been implicated in cell extrusion in simple epithelia^26^. Migrating keratinocytes typically display pronounced posterior enrichment of α6 integrin^51^, which was absent here (**ED Figure 7b**). Together with the inward orientation of traction forces, these findings argue against directed neighbor migration as the source of the observed flows. Instead, they support the model prediction that convergent tissue motion emerges from a differentiation-induced, three-dimensional redistribution of active stresses.

**Figure 6:**
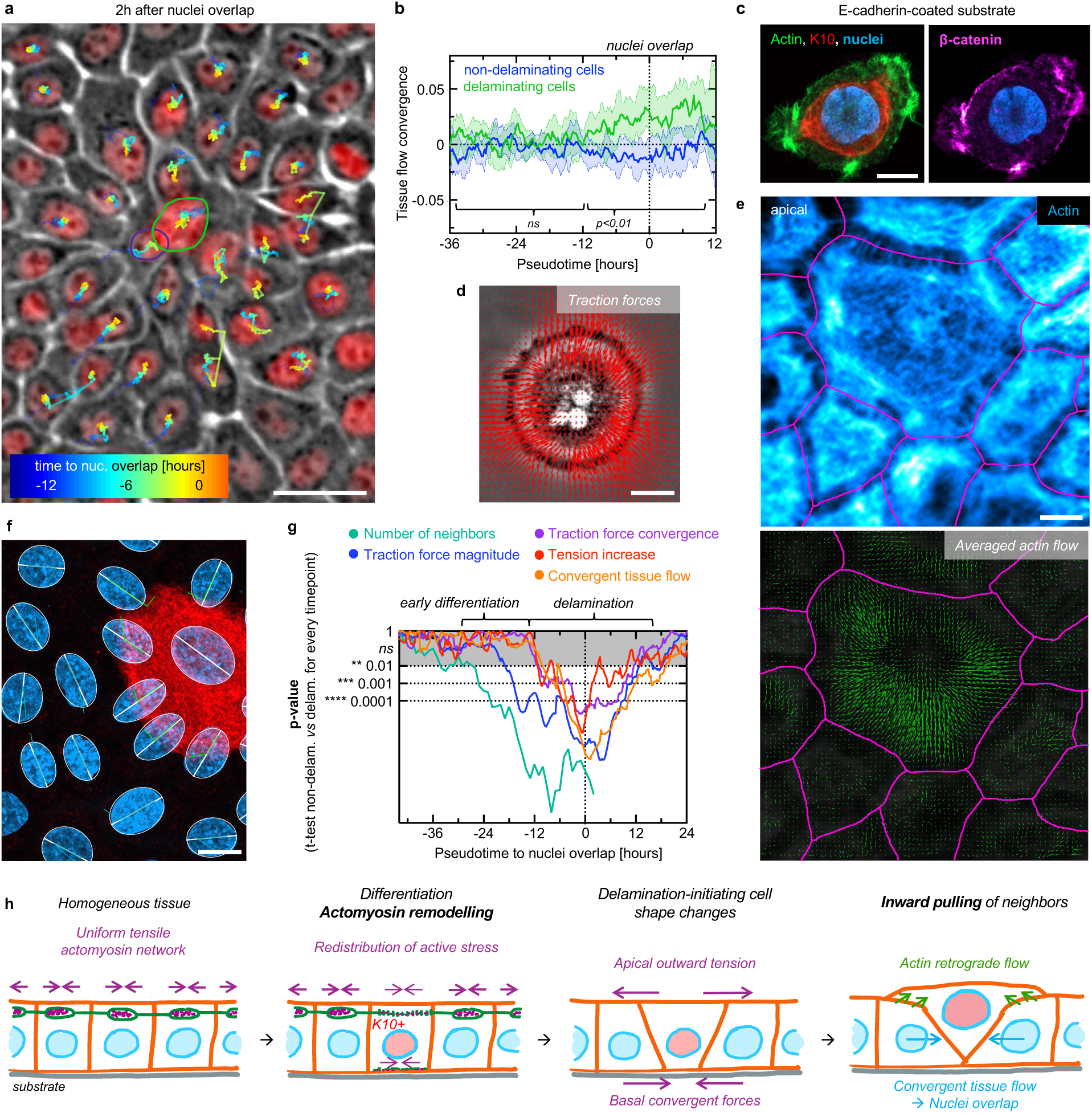
Convergent tissue flows close the basal layer below delaminating cells. **a** Phase contrast image of a delaminating cell 2 hours after nuclei overlap. Nuclei were tracked over 16 hours. Their trajectories are color-coded based on time to nuclei overlap from blue (early) to orange (late). **b** Local tissue flow orientation (convergence) as a function of time to nuclei overlap. Median +/− 95% CI of n=67 individually tracked cells from N=3 independent experiments. p-value from continuous t-test; brackets indicate timeframe where p<0.01 or not significant (ns). **c** Confocal images of a differentiated cell plated on an E-cadherin-coated substrate showing nuclei (blue), actin (green), K10 (red) and beta-catenin (magenta). **d** Traction force field generated on E-cadherin. **e** Top: representative image of the apical actin (blue). Bottom: averaged actin flow during a 15-hour period. Cell outlines are indicated in magenta. **f** Nuclei orientation close to a K10+ cell. Nuclear ellipses and long axis in white, angles relative to the apical interface of the delaminating cell. **g** Temporal comparison of the difference between delaminating and non-delaminating cells for several mechanical signatures. P-values were computed for every timepoint relative to the nuclei overlap for n=67 events from N=3 independent experiments. **h** Schematic of how differentiation-induced actomyosin remodeling and force exertion generate a delamination-promoting environment. Scale bars 10 µm (**c**, **d**, **e**, **f**); 25 µm (**a**).

### Delaminating cells pull and reorient their neighbors through a retrograde actin flow

To identify the origin of the convergent flows, we tested whether neighbor cells could be actively pulled inward by the delaminating cell. To isolate this mechanism, we coated TFM substrates with E-cadherin and plated individual differentiated cells, allowing them to engage the substrate cadherin-mediated cell-cell adhesions (**Figure 6c**). Strikingly, these cells generated a radially inward-pointing force pattern (**Figure 6d**), that closely resembled the aster-like traction force defect observed during delamination. Consequently, analysis of bead displacements revealed convergent deformation of the underlying substrate (**ED Figure 7c**). These results demonstrate that differentiating cells are intrinsically capable of transmitting inward-directed forces through E-cadherin-based adhesions. They support a model in which the delaminating cell actively pulls its immediate neighbors, providing a direct mechanical origin for the convergent tissue flows observed in the intact epithelium.

Such inward-pulling forces can be generated through retrograde actin flows^52^, which were previously recorded in single keratinocytes^30^. We observed such retrograde actin flows within apically expanded cells (**Figure 6e, ED Figure 8a***, Supplementary Movie 10*), which could pull the neighbor cells inward and even orient and deform them. Indeed, quantifying neighbor cells nuclei’s angles relative to the apical interface of differentiating cells (**Figure 6f**) revealed a predominantly perpendicular orientation in neighbors but not in cells further away (**ED Figure 8b**), supporting pulling forces exerted by delaminating cells. Neighbor cells also underwent pronounced three-dimensional shape and polarity changes. Compared with distant cells, they displayed a larger offset between their apical and basal areas (**ED Figure 8c**); their apical domains shifted away from the delaminating cell, whereas their basal domains displaced towards it (**ED Figure 8d**). This geometric polarization was accompanied by a reorganization of intracellular polarity. In distant cells, the Golgi apparatus remained close to the nucleus without a preferred orientation, whereas in immediate neighbors it was consistently displaced away from the delaminating cell (**ED Figure 8e,f**). Together, these findings support a mechanism in which differentiation-associated actin retrograde flows enable the delaminating cell to pull on its neighbors. This inward force not only drives convergent tissue motion, but also imposes local orientation and intracellular polarity, revealing the delaminating cell as an active mechanical organizer of its surrounding tissue.

### Temporal evolution of the mechanical signatures of cell delamination

To integrate the cell- and tissue scale mechanical changes that culminate in delamination, we mapped their temporal sequence by comparing delaminating and non-delaminating cells at each timepoint (**Figure 6g**). The earliest significant difference was an increased neighbor number around one day before the nuclei overlap, consistent with the progressive enlargement of differentiating cells. This was followed by an increase in traction-force magnitude, which we attribute to enhanced force generation associated with elevated phospho-myosin levels. Together, these early changes mark the emergence of differentiation-driven mechanical heterogeneity within the tissue. A major transition occurred approximately 12 hours before nuclei overlap, when several mechanical signatures appeared nearly simultaneously. Local tensile stresses increased; traction forces became convergent and organized into a +1 topological force defect, and the surrounding tissue developed convergent flows. The signatures remained elevated for approximately one day and returned to the levels observed around non-delaminating cells 10-20 hours after the nuclei overlap (**Figure 6g**). Because this coordinated mechanical state is reproduced by our hydrodynamic model, we propose that it drives delamination-initiating shape changes. Together, delamination is not an acute event, but a prolonged, mechanically coordinated process unfolding over at least 1.5 days, in close agreement with its timescale *in vivo*^18^.

## Discussion

Here, we address a longstanding question in epidermal biology: what initiates delamination in a dense and homeostatic tissue? By combining quantitative biophysical experiments with a hydrodynamic theoretical model, we show that delamination emerges from differentiation-induced mechanical heterogeneity that reorganizes forces across the plane and the height of the epithelium. We directly measure the forces and stresses during delamination and identify apicobasal actomyosin remodeling as a key event that redistributes active stress within differentiating cells. Our model shows that this vertical redistribution is sufficient to generate outward-directed forces apically and inward-directed forces basally, thereby reshaping the cell-cell interface and driving the characteristic cell-shape changes and convergent tissue flows that precede delamination. The model predictions are in line with experimental observations of increased tissue tension generated on the apical side, the emergence of an inward-pointing +1 traction force defect on the basal side and the development of convergent flows in the surrounding tissue (**Figure 6h**).

A central feature of this transition is the extensive reorganization of the actomyosin cytoskeleton. Apical stress fibers are replaced by orthoradial actin arcs, accompanied by pronounced retrograde actin flow. This reorganization precedes the formation of the honeycomb-like actin network that develops in suprabasal cells after several days of differentiation induction and resembles actin organizations observed *in vivo*^11,15,29^. Our data therefore suggests that basal keratinocytes transiently enhance retrograde actin flows as part of their differentiation program and use it to transmit basal inward-directed forces to neighbor cells. In this view, delamination is neither a purely cell-autonomous process nor a passive response imposed by the surrounding tissue. It is initiated within the differentiating cell through actomyosin remodeling but completed through the mechanical response of its neighbors. By integrating tissue stresses over full epithelial height^42^, we detected a net increase in tension during delamination. Importantly, our model resolves the three-dimensional stress distribution underlying this integrated measurement and predicts that active stresses change sign along the apicobasal axis, producing basal compression together with apical tension. Thus, although apical stresses dominate the overall mechanical state of the tissue here, it coexists with a localized basal compressive component. This reconciles our findings with basal compression measured in more matured tissues^15^, and supports a contribution of increasing basal compression in initiating delamination, as hypothesized previously^23^. More broadly, these results emphasize that measurements restricted to a single plane may obscure mechanically opposing contributions distributed across epithelial height. Delamination should therefore be understood as the outcome of a three-dimensional stress imbalance rather than a response to planar tension or compression.

Substrate stiffness can modulate traction forces^53,54^, and consequently the friction length that governs tissue-force transmission^55^. Basement membrane stiffness has been reported to increase by approximately 2.5-fold during development^56^ and to decrease by roughly 4-fold in aging-related phenotypes^57^. In our model, varying the friction length over a 5-fold range did not qualitatively alter the predicted delamination-associated shape changes, suggesting that physiological variations in basement-membrane stiffness are unlikely to abolish the core mechanism identified here. Nevertheless, the model predicts that friction influences the timescale over which delamination unfolds. Changes in matrix mechanics may thus regulate the rate, rather than the fundamental geometry of cell exit and could thereby modulate the pace of epidermal stratification or renewal. Testing this prediction will require direct manipulation of basement-membrane mechanics in physiologically relevant systems.

Our experiments capture delamination during formation of the first suprabasal layer whereas *in vivo* basal cells usually delaminate into an already established stratified epithelium. The mechanism proposed here does not rely on potential contributions from suprabasal layers, as it only requires the apicobasal redistribution of active stresses within the basal cell layer. In principle, reducing apical in-plane tension in a basal cell embedded within a multilayered tissue could therefore generate similar shape changes. The existence of such an apical tensile environment in the basal epidermal layer is plausible, as actomyosin accumulates at the interface between basal and suprabasal layers during development^41,56^, and in adult mice epidermis^11,58^. However, whether these structures are generated apically by basal cells or basally by suprabasal cells remains unresolved. Thus, further research needs to address to which extend the delamination mechanism proposed here can be applied to stratified tissues, also because mechanical interaction between the layers may change it, as such forces were shown to affect basal cell fate^58^. Despite these open questions, the structural and regulatory principles uncovered here may extend beyond the epidermis. Several multilayered epithelia like thymus, esophagus, bladder or prostate share aspects of epithelial architecture and differentiation programs with the epidermis^59,60^. Our results therefore raise the possibility that differentiation-driven redistribution of active stress represents a more general mechanism for coordinating cell exit in self-renewing tissues. More broadly, our study identifies differentiating cells not as passive products of tissue organization, but as active mechanical organizers that reshape their neighborhood and create the local force landscape required for their own delamination.

## Methods

### Cell culture

Human N/TERT-1 keratinocytes, isolated from newborn foreskin, were originally established by James Rheinwald (Harvard University)^27^, who kindly allowed its use. Cells were grown in growth medium (CnT-Prime, CellnTech) supplemented with 1% penicillin/streptomycin. The growth medium contains negligible amounts of calcium (0.07mM). Cells were incubated at 37°C and 5% CO^2^. Cells were passaged using trypsin. Prior to cell seeding, substrates were incubated with 0.04 µg/ml Collagen type-I (Biochrom L7213) in PBS for 30 min at room temperature, and rinsed 2x with PBS. For differentiation experiments, cells were seeded into Collagen type-1 coated glass-bottom fluorodish in 2 ml growth medium to reach confluence. To induce differentiation, cells were switched to differentiation medium (CnT-Prime-3D, CellnTech), which contains high calcium (1.2 mM) and was supplemented with 1% penicillin/streptomycin. The differentiation medium was replaced every two days.

### Indirect immunostaining

Cells were rinsed with warm PBS. Fixation was carried out in 4% formaldehyde for 10 min at room temperature and at 37°C for tubulin staining. Cells were permeabilized using 1% Triton X-100 in PBS for 15 min followed by 3 x 5 min washing in PBS. Samples were blocked with 4% BSA in PBS for 1 h at room temperature. Unless otherwise stated, all following primary antibodies were diluted 1:200 in blocking solution (4% BSA in PBS) and incubated for 1-2 h at room temperature or over night at 4°C.

- anti Keratin 10 (K10) rabbit antibody (catalog no 905404, Biolegend)
- anti beta-catenin mouse antibody (catalog no 610156, BD Biosciences)
- anti phospho-myosin light chain 2 (pMLC2) mouse antibody (catalog no 3675, Cell Signaling) – diluted 1:300
- anti Desmoglein 1/2 mouse antibody (catalog no 61002, Progen)
- anti E-cadherin mouse antibody (catalog no 610181, BD Biosciences)
- anti GM130 mouse antibody (catalog no 610822, BD Biosciences)
- anti alpha-tubulin (TUBA1) mouse antibody (catalog no T9026, Sigma)
- anti alpha-6 integrin rat antibody (catalog no MAB13501, RCD Systems)

The samples were washed 3 x 5 min with PBS and incubated with anti-rabbit immunoglobulin (catalog no A31573, Life Technologies) or an anti-mouse immunoglobulin (catalog no A31571, Life Technologies) antibodies conjugated to Alexa Fluor 647 or Alexa Fluor 568 diluted 1:200 in blocking solution for 1-2 h at room temperature. Subsequently, samples were washed 3 x 5 min with PBS. The actin cytoskeleton was visualized using Phalloidin-Alexa Fluor 488 or Alexa Fluor 555 (catalog no A12380, Life Technologies) diluted 1:200 in blocking solution and the nuclei using Hoechst 33342 (catalog no 62249, Thermo Fisher) diluted 1:2000 in PBS for 45 min at room temperature.

### Confocal microscopy, data visualization and image analysis

Fixed samples were imaged on a laser scanning confocal microscope (Zeiss LSM 980, Germany), equipped with a 63x oil objective and an Airyscan 2 module. All images or Z-stacks were acquired in Airyscan mode without further averaging, and an automated deconvolution was performed within the microscope software (Zeiss ZEN blue). All images were visualized using Fiji^61^ and brightness and contrast were adjusted. For Z-stacks, maximum intensity projections, sum projections, height projections or side views were generated.

#### Measurements of area, number of neighbors and approximation of cell contractility

Z-stacks covering the height of the cells were acquired one day after differentiation induction for pMLC2, actin, nuclei and K10 staining. The following cell populations were compared: Differentiating cells, identified by K10 expression, 3 randomly selected direct neighbors, and 4 randomly selected, non-differentiated cells more than 1 row away from the differentiating cells. For all cells, we manually counted the number of neighbors. Nuclear areas were measured manually based on the nuclear staining and cell areas in the apical or the basal plane based on actin staining. The ratio between basal and apical areas were calculated for all cells. The total amount of phospho-myosin as a proxy for cell contractility was assessed through a sum-projection of the whole Z-stack. Phospho-myosin levels were measured for all cells in the areas segmented based on actin staining. To compare across experiments, levels were normalized by the average of the random cells > 1 row away from the differentiating cells.

#### Quantification of actomyosin reorganization in delaminating cells

Z-stacks were acquired after 2 days following differentiation induction as described above and images were analyzed in Fiji. K10+ cells were selected. To compare their apical and basal actomyosin levels with the neighbors, apical and basal areas were segmented manually in the appropriate z-plane. The mean fluorescence intensity per unit area were measured and normalized to the average intensities from 3 randomly selected neighbor cells. To quantify the shift in vertical (apical-basal) distribution, the ratio between the mean fluorescence intensities per unit area on the apical and basal side were computed. To assess intracellular, horizontal actomyosin remodeling, line plots with a width of 5 µm were drawn spanning the apical or basal planes of delaminating cells and their neighbors. The intensities were normalized to the maximal value and rescaled to the maximum length, which varied between delaminating and non-delaminating cells.

#### Quantification of cell polarity

Z-stacks covering the height of the cells were acquired after 1 and 2 days of differentiation induction for E-cadherin, actin, GM130, K10 and nuclei stainings. To investigate apical-basal area shifts relative to the delaminating cell, the centers of mass of the apical and the basal area were measured and the offset was calculated. Then, the center of mass of the delaminating cell was determined and the distances between delaminating cell and apical or basal area were calculated. If the distance between the apical area and the delaminating cell was greater than the distance between the basal area, the apical-basal shift was determined positive.

### Bulk stiffness measurements

Nanoindentation was used to measure the stiffness of cell monolayers grown on collagen-coated glass, in all cases after confluency; before differentiation induction and 1, 2 or 3 days afterwards. The indenter (Chiaro Nanoindenter, Optics 11 life, Netherlands) was connected to an epifluorescence microscope equipped with a 10x objective to visualize the indentation position. A soft cantilever (k = 0.015 N/m) with a small tip (radius = 3 µm) was calibrated on glass before the measurement according to the manufacturers protocol. Before every measurement, the distance of the probe to the surface of the sample was determined automatically, and the probe was placed 5 µm above the surface. To measure the stiffness, the cantilever was pushed down 10 µm, held there for 2 sec and retracted afterwards. The matrix scan function was used with a typical step size of 25 µm. Measurements were conducted at random positions. To determine the elastic modulus, the loading curve was analyzed using the built-in software (DataViewer V2, Optics 11 life). The analysis is based on the Hertz model (Hertzian contact), which assumes a linear elastic response of the sample. The single fit method was used with a maximal load (P_max_) of 90% and a Poisson’s ratio of 0.5. Loading curves, which started in contact due to a failed determination of the surface or curves where the surface was not found at all were excluded. Quality control for the fitted loading curves was performed with a cut-off of 0.97.

### Video microscopy

#### Substrate preparation

15 kPa soft silicone substrates for TFM were prepared as described previously ^25,42^. In brief, CY52-276A and CY52-276B polydimethylsiloxane (Dow Corning, Toray) were mixed in a weight ratio of 1:1 and then poured on glass bottom imaging dishes (Fluorodish, WPI) to obtain a layer of thickness around 100 µm and equilibrated for 30 min. The substrates were then cured at 80°C for 2 hours. Prior to seeding of the beads, the surface was silanized using a solution of 5% APTES ((3-Aminopropyl)triethoxysilane, Sigma-Aldrich) diluted at 10% in absolute ethanol for 10 min, then washed with absolute ethanol 3 times, before being dried at 80°C for about 10 min. 200 nm red carboxylated fluorescent beads (FluoSpheres, Invitrogen) were diluted within a 2:1000 ratio in water, and subjected to an ultrasonic bath for 10 min. The beads solution was then filtered using a 0.22 µm filter and incubated on the substrates for 15 min, protected from light. The dishes were finally washed with water 3 times and dried at 80°C for 3 minutes.

#### Surface coating

Prior cell seeding, these substrates were coated with 50 µg/mL fibronectin (Sigma) for 1 hour and washed 3 times with PBS. Other substrates were coated with 2.5 µg/100 µl recombinant human E-cadherin Fc (catalog no. 10204-H02H, SinoBiological) diluted in PBS from a 250 µg/ml stock solution. E-cadherin Fc was incubated for 1 hour at room temperature. Substrates were washed 3 times with PBS, treated with Pluronix for 1 hour, and washed 3 times with PBS.

#### Time-lapse microscopy

Cells were seeded on collagen-coated glass and for TFM experiments on fibronectin-coated TFM substrates in growth medium. Once confluence was reached, differentiation was induced and time-lapse microscopy was carried out using Nikon Biostation microscopes equipped with a 10x phase contrast objective (for TFM) or a Nikon Spinning Disk (CSU W1, equipped with a 20x air objective). To fluorescently label nuclei, SPY650-DNA (1:750, Spirochrome) was added 1-2 hours before data acquisition or together with the differentiation medium. To label actin, SPY555-FastAct_X (1:1000, Spirochrome) was added 3-4 hours before data acquisition. To capture the full dynamics of differentiation, medium switches or changes were performed directly on the microscope, by carefully removing the old medium and adding new differentiation medium. After completion of the TFM experiment, cells were removed by adding SDS and the reference bead image was taken 1 hour after SDS treatment. To investigate forces exerted through E-cadherin contacts of differentiated cells, cells were pre-differentiated for three days as described above. Cells were trypsinized and seeded onto E-cadherin Fc-coated TFM substrates in differentiation medium to obtain single cells. Non-adherent cells were washed away after 3 hours of incubation. The sample was then processed as fibronectin-coated samples as described above to measure traction forces.

### Laser ablation

Confluent cultures, after 1 day of differentiation induction, were incubated with SPY555-FastAct_X (1:1000, Spirochrome). Laser ablation was performed on a Nikon CSU-X1 (Yokagawa) spinning disc microscope with a fluorescence recovery after photobleaching module (iLAS2, GATACA) and a 40x/1.2 water-immersion objective. Lines spanning 3-4 cells were ablated by focusing a UV laser (355 nm, pulse duration 3-5 ns, laser power 450 mW) for 1 sec. Samples were imaged 1 minutes before ablation and immediately after with a framerate of 1 sec. The recoil velocity was measured manually within the first 5 seconds following ablation by segmenting the holes over time and computing the edge displacement speed of the ablated regions.

### Video microscopy analysis

#### Tracking of delaminating cells

Fluorescent nuclei data based on SpyDNA staining was pre-processed, depending on its initial quality using Fiji. First, a background subtraction was performed using 100 pixel rolling ball radius and a sliding paraboloid. The image was smoothed with a 2×2 pixel gaussian filter. A second background subtraction was performed if necessary, and the image was smoothed using two times a 2×2 pixel median filter. If further necessary local contrast was enhanced using CLAHE. An overlay of the phase-contrast and the processed fluorescent nuclei channel was generated and delaminating cells were tracked semi-automatically over the whole movie using the manual tracking pipeline of TrackMate v7^62^. The quality threshold was set to 0.3 and the distance tolerance was set to 1. The spot radius was adjusted to fit the nuclear size for each cell. Delaminating cells were identified manually in later frames of the movie where the overlap of two nuclei became visible. The beginning of an overlap of the nuclei of delaminating cell and a neighboring cell marked the reference timepoint of delamination and was determined manually and noted in “Tracks”. For each delamination, a cell which was not delaminating, but divided at least once during the movie was tracked as a control. To correct for tissue-scale changes (i.e. phase transitions) depending on differentiation time, the reference timepoint for the control cell was set identical to the reference timepoint of delamination. CSV tables of “Tracks” and “Spots” were exported from TrackMate and further processed using MATLAB. Overlays of the tracking and the input data were exported as movies. The number of delamination events was extracted as a function of time after differentiation induction.

#### Velocity measurements

Particle Image Velocimetry (PIV) tissue flow fields were generated for the whole field of view using PIVlab ^63^, a toolbox developed in MATLAB, comparing consecutive frames with an interrogation window of 16×16 pixels (∼10×10 µm) and a 50% overlap. Cell movements within these windows were averaged across the whole field of view and over time to compute the tissue speed.

#### Traction force measurements

The bead images obtained during the time-lapse microscopy were merged with corresponding reference bead images taken after SDS treatment. The resulting stack of images was pre-processed using the Image Stabilizer plug-in in ImageJ and the illumination was corrected to remove background noise. Displacement field of beads was obtained using PIVlab^63^ with an interrogation window of 16×16 pixels and an overlap of 50%. Bead displacements were then correlated to a traction force field using Fourier transform traction cytometry (FTTC), the known substrate stiffness of 15 kPa, and a regularization parameter of 9×10^−9^.

#### Stress measurements

From traction force fields, we inferred the stress tensor everywhere in the tissue using Bayesian inversion stress microscopy ^42,43^ with a regularization parameter **Λ** = 10^−6^. Isotropic stress was calculated as half of the trace of the stress tensor. To generate heatmaps of isotropic stress and traction force magnitude, a smoothing was applicated through linear interpolation.

#### Local analysis of mechanical properties and local tissue flows direction

Local readout of mechanical properties was done using MATLAB. Based on the tracking data (“Spots”), a small (25 x 25 µm) window was placed around each spot at each timepoint, for all detected events. Then, traction force magnitudes, the traction force divergence and isotropic stresses were averaged in each window, providing the frame-resolved local data, from start to finish of the movie. Then, the data was aligned based on the delamination timepoint: the frame number was shifted for each event that frame zero corresponds to the timepoint which was previously manually defined as the nuclei overlap. This resulted in a shifted dataset showing the pseudo-time around the delamination timepoint. To assess tissue flow directions, 50 x 50 µm windows were used to locally read out PIV-based tissue flows. Local vectors were normalized by their length to quantify the divergence of the local cell movement through time. As values became predominantly negative for delaminating cells, we inverted the divergence and reported the convergence for traction forces and tissue flows. All data was exported as time-resolved data for single events with the timepoint zero being the detected nuclear overlap. The data was smoothed by performing two times a rolling average over 9 frames, corresponding to +/−1 hour around each timepoint. Data was visualized using GraphPad Prism showing the median and the 95% confidence interval.

#### Averaged traction force fields around delaminations and comparison with aster

Using a custom MATLAB code, traction forces vectors were overlayed and averaged in a window of 250 x 250 µm surrounding the delaminating and non-delaminating cells for all available data. This was done starting 16 hours before the nuclei overlap and 6 hours afterwards. Vectors were smoothed using a rolling average of 1 hour, or 5 frames. Averaging led to a readout of a traction force increase, as vectors of the same length (force magnitude), but opposite orientation annihilate each other and result in a zero-traction force readout. These forces were averaged radially to obtain the tangential component and fitted with an exponential decay function in GraphPad Prism. An aster-shaped +1 topological defect was simulated in MATLAB with the same vector spacing as in the experimental data. The averaged experimental traction force vectors were then compared to the aster and a correlation score between −1 (vectors in opposite direction) and 1 (vectors overlap perfectly) was computed for each vector, resulting in a time-resolved correlation map. Starting from the center, the correlation score was first radially averaged for all timepoints and then averaged for 2-hour intervals to obtain the time-and space-resolved correlation between experimental data and the aster.

### Statistical tests

To assess the development of statistical significances as a function of time, two-sided t-test were conducted between the datasets of the delaminating and non-delaminating cells for every timepoint. The obtained p-value was visualized using GraphPad Prism (V10). All further statistical tests were directly performed in GraphPad Prism (V10).

## Acknowledgments

We thank the members of the “Cell Adhesion and Mechanics” team for helpful discussions, particularly Marc-Antoine Fardin, Philippe Marcq for feedback on BISM, Danijela Vignjevic and Nicolas Borghi. We thank James Rheinwald for allowing to use N/TERT-1 keratinocytes. We acknowledge the ImagoSeine core facility of the Institut Jacques Monod, member of the France BioImaging infrastructure (https://ror.org/01y7vt929) supported by the French National ResearchAgency (ANR-24-INBS-0005 FBI BIOGEN) and GIS-IBiSA and the support of the Fondation Bettencourt-Schueller.

## Author contributions

A.S., R.V., B.L., C.N., R.M.M. conceived the study. A.S. designed all experiments and performed and analyzed them together with C.D. J.O.B. developed the theoretical model. W.S. performed numerical simulations. L.A. and Y.S. developed analysis codes. R.D. contributed to establishing protocols. F.W. performed laser ablation experiments. A.S., R.V., B.L., C.N., R.M.M. wrote the manuscript and all authors provided feedback on it.

---

**Extended Data Figure 1:**
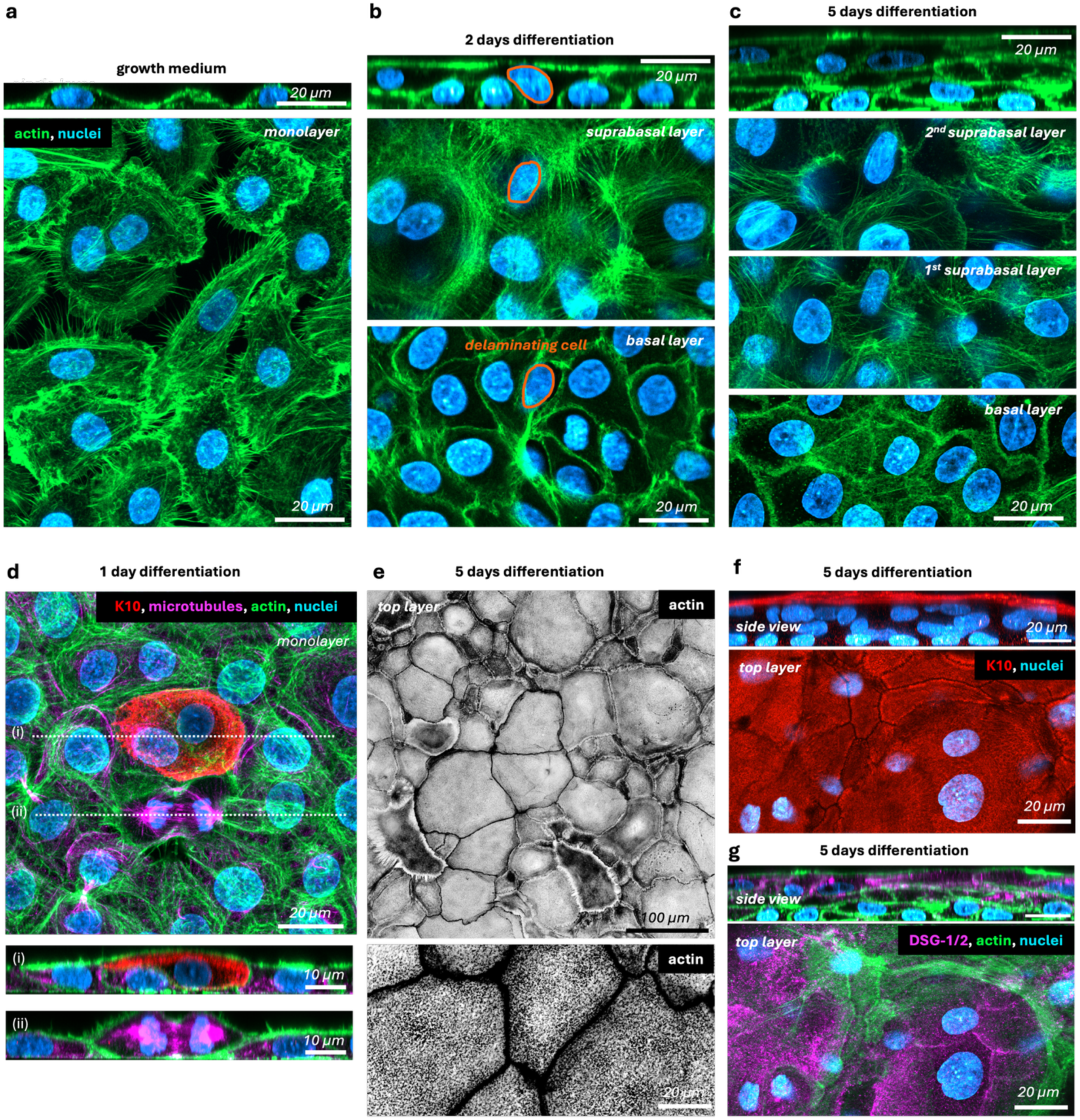
Organotypic multilayer formation of N/TERT-1 keratinocytes. **a** Vertical- and horizontal views of a N/TERT-1 keratinocyte culture before the induction of differentiation. Actin in green, nuclei in blue. **b** Vertical and horizontal views of the basal and emerging suprabasal layer 2 days after differentiation induction. **c** Vertical and horizontal views of the basal, 1^st^ and 2^nd^ suprabasal layer 5 days after differentiation induction. **d** Horizontal view of the monolayer 1 day after differentiation induction. K10 in red, microtubules in magenta. Side views of a delaminating (i) and a dividing cell (ii), illustrating how the suprabasal layer is generated independent of cell division. **e** Top: horizontal view of the outermost layer 5 days after differentiation induction. Actin in inverted greyscale. Bottom: Higher magnification and adjusted contrast to visualize the actin cortex. **f** Vertical- and horizontal view of K10 (red) in the outermost layer 5 days after differentiation induction. **g** Vertical- and horizontal view of Desmosomes (magenta) in the outermost layer 5 days after differentiation induction.

**Extended Data Figure 2:**
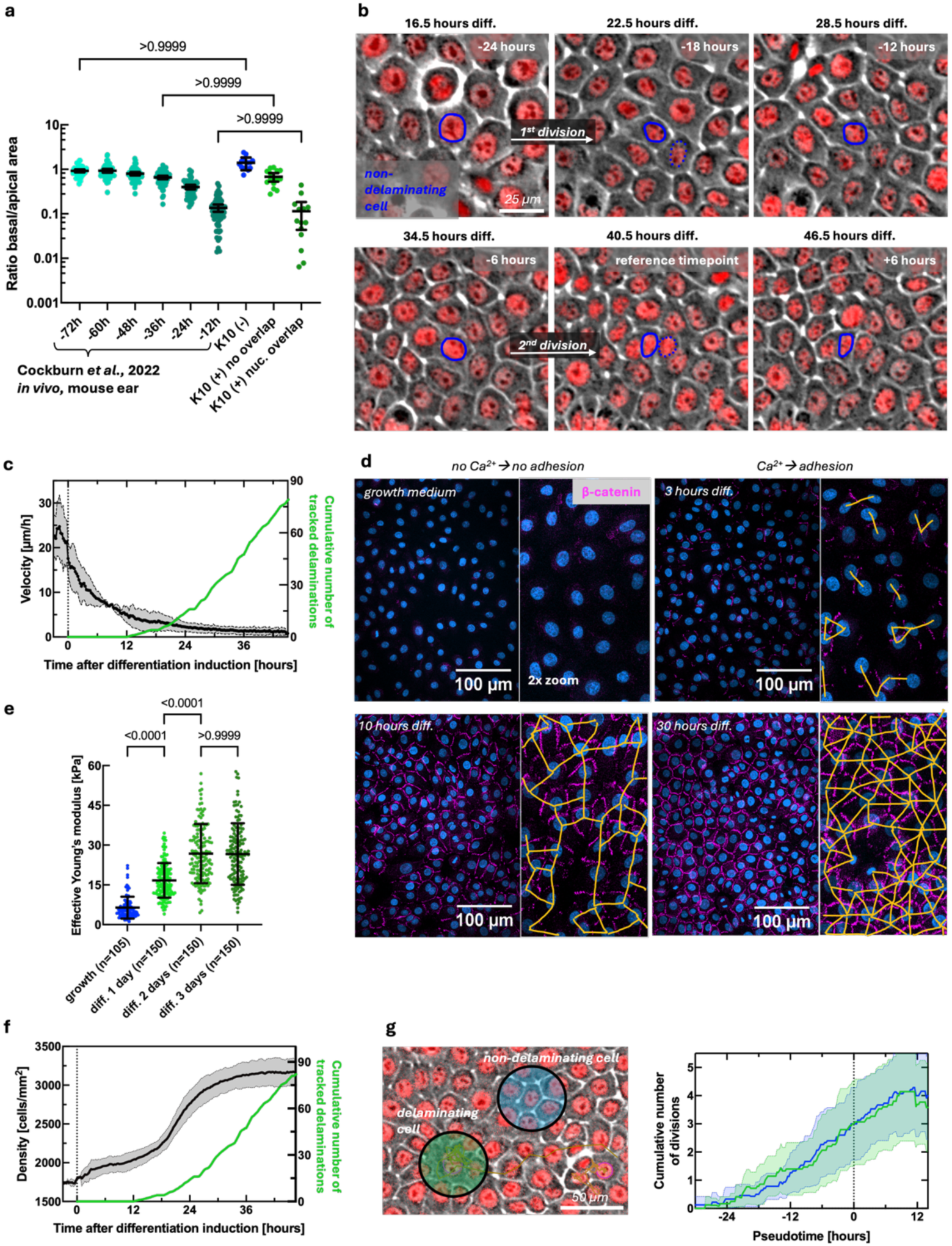
Global physical changes precede delamination. **a** Comparison of the basal-to-apical area ratio during the progression of homeostatic delaminations in the mouse ear reported by Cockburn *et al.*, Nature Cell Biology, 2022 and the ones reported here (data from Figure 1i). **b** Example phase contrast images of a tracked non-delaminating but proliferative cell. After cell divisions, tracking was continued for one random daughter cell. The reference timepoint corresponds to nuclei overlap for a delaminating cell (see Figure 1). **c** Tissue velocity measured by PIV as a function of time after differentiation induction. Green: Cumulative number of tracked delaminations. **d** Representative confocal images of the tissue following differentiation induction. Beta-catenin in magenta. Yellow lines show cell-cell adhesion, which eventually spans the whole tissue. **e** Effective Young’s modulus measured by indentation at different timepoints relative to differentiation induction. Every datapoint shows 1 measurement from N=2 independent experiments. P-values from multiple ANOVA tests. **f** Tissue density as a function of time after differentiation induction. Green: Cumulative number of tracked delaminations. **g** *Left*: Example image of the local neighborhoods of delaminating and non-delaminating cells in which cell divisions were counted. *Right*: Cumulative number of divisions relative to the nuclei overlap at t=0. For all plots, means +/− SD.

**Extended Data Figure 3:**
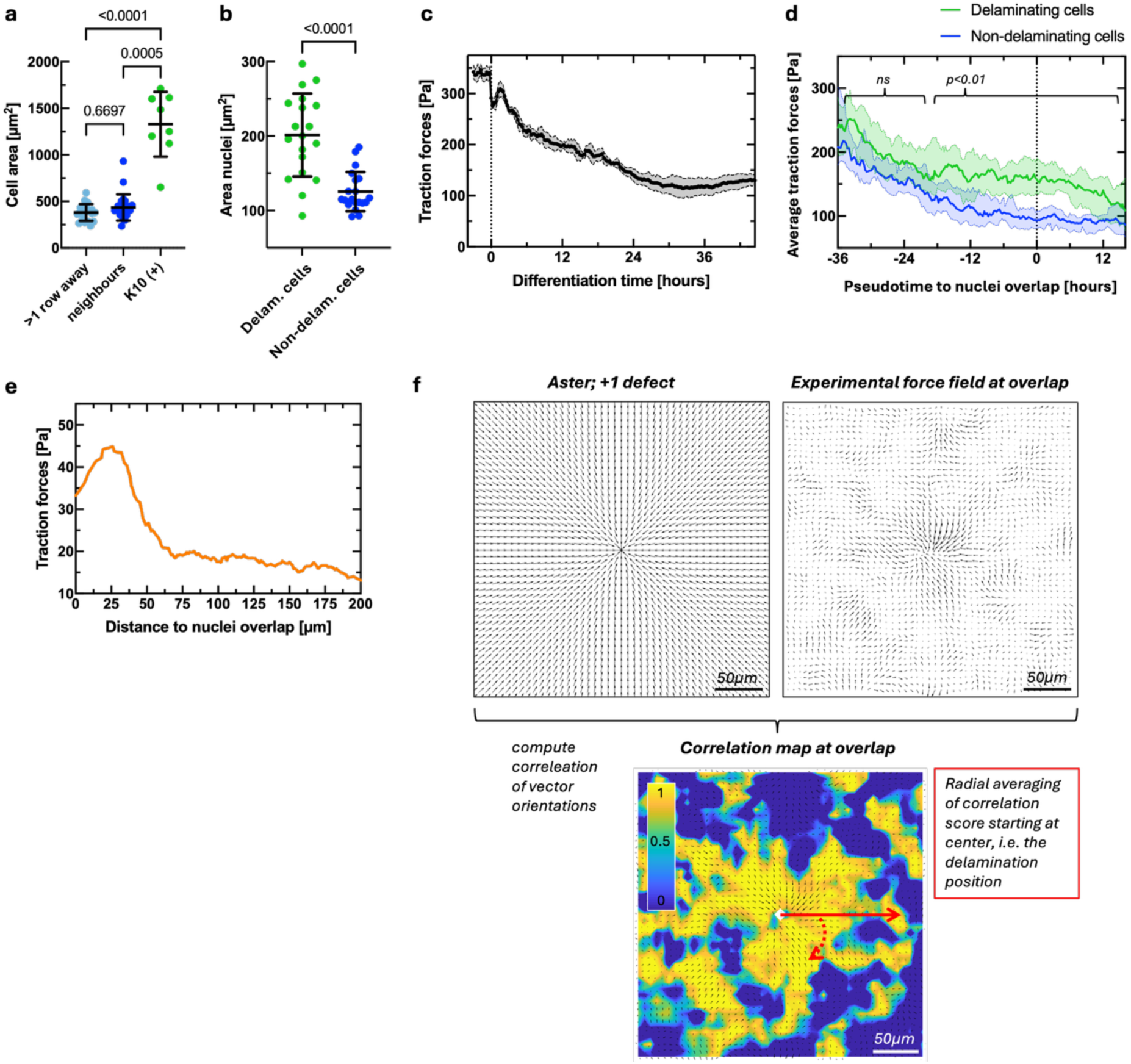
Emergence of a +1 defect. **a** Cell area of non-differentiated and differentiated cells 1 day after differentiation induction measured in fixed images. **b** Nuclei area of delaminating and non-delaminating cells 24 hours before the nuclei overlap measured in movies. **c** Global traction forces following differentiation induction **d** Local traction force magnitude around delaminating and non-delaminating cells relative to the nuclei overlap at t-0. P-values from continuous t-test. **e** Radial distribution of the oriented traction forces at nuclei overlaps. **f** Pipeline to compute the correlation between an aster-shaped +1 topological defect (*left*) and the experimental averaged traction force field (*middle*), resulting in a correlation map (*right*). To compute correlation scores at a given timepoint, the correlation was radially averaged (*red arrow*) starting at the defect center, which is the position of the delaminating cell’s nucleus.

**Extended Data Figure 4:**
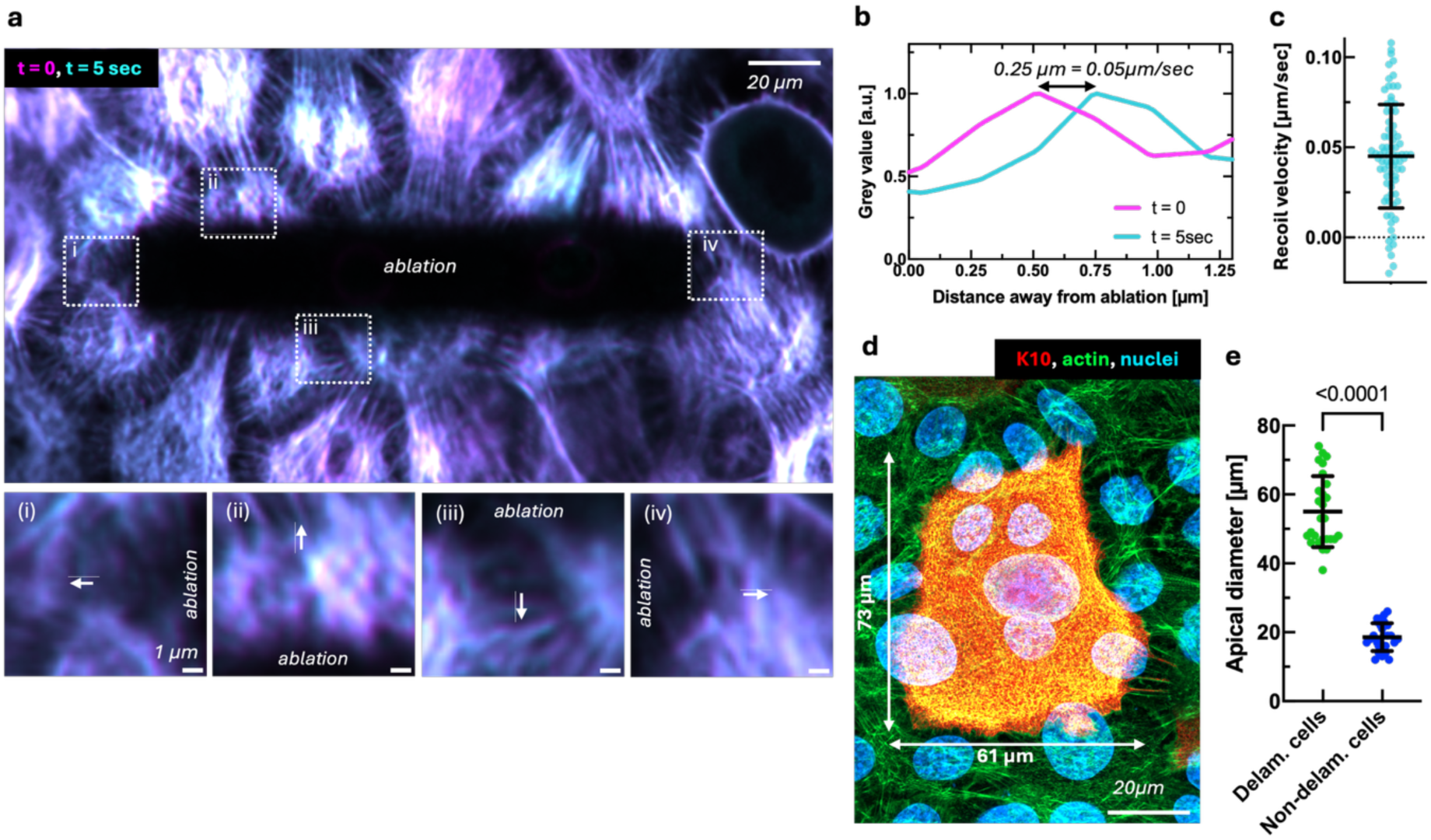
Laser ablation reveals tissue-level tension. **a** Representative overlayed image of a laser-ablated area directly after the ablation (t=0, magenta) and 5 seconds later (blue). Zoom-ins show an isotropic motion of non-ablated actin away from the ablated area. **b** Example quantification of the recoil velocity. The spatial shift in actin intensity peaks was measured after 5 seconds, and the obtained distance was converted into a speed. **c** Recoil velocity of the actin cytoskeleton during the first 5 seconds following laser ablation 1.5 days after differentiation induction. Each datapoint shows one measurement from N=2 independent experiments. **d** Confocal image of the apical expansion of a differentiating cell at nuclei overlap (K10: red, actin: green, nuclei: blue) on top of its neighbors. **e** Quantification of the apical diameters of delaminating and non-delaminating cells 2 days after differentiation induction.

**Extended Data Figure 5:**
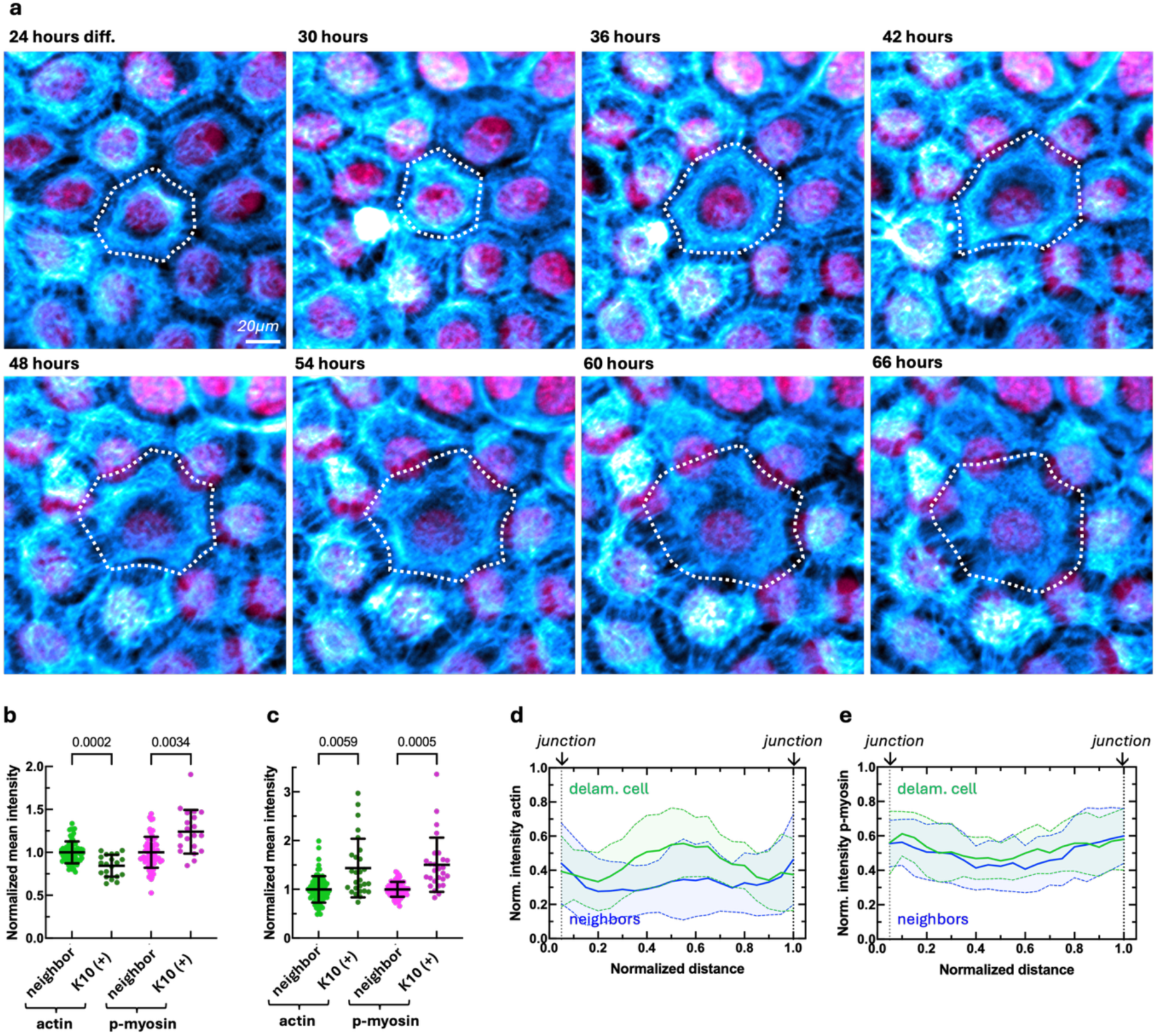
Actomyosin remodeling. **a** Snapshots of the apical actin (blue) in an apically expanding cell over 42 hours. Cell outlines in white. Nuclei in pink. **b** Comparison of the mean actomyosin intensity per unit area in the apical plane, normalized to the mean value of the neighbor cells. **c** Comparison of the mean actomyosin intensity per unit area in the basal plane, normalized to the mean of the neighbor cells. **d** Horizontal distribution of actin and **e** phospho-myosin intensity across the basal plane of delaminating (green) and non-delaminating cells. Intensity and distance were normalized to the maximum value.

**Extended Data Figure 6:**
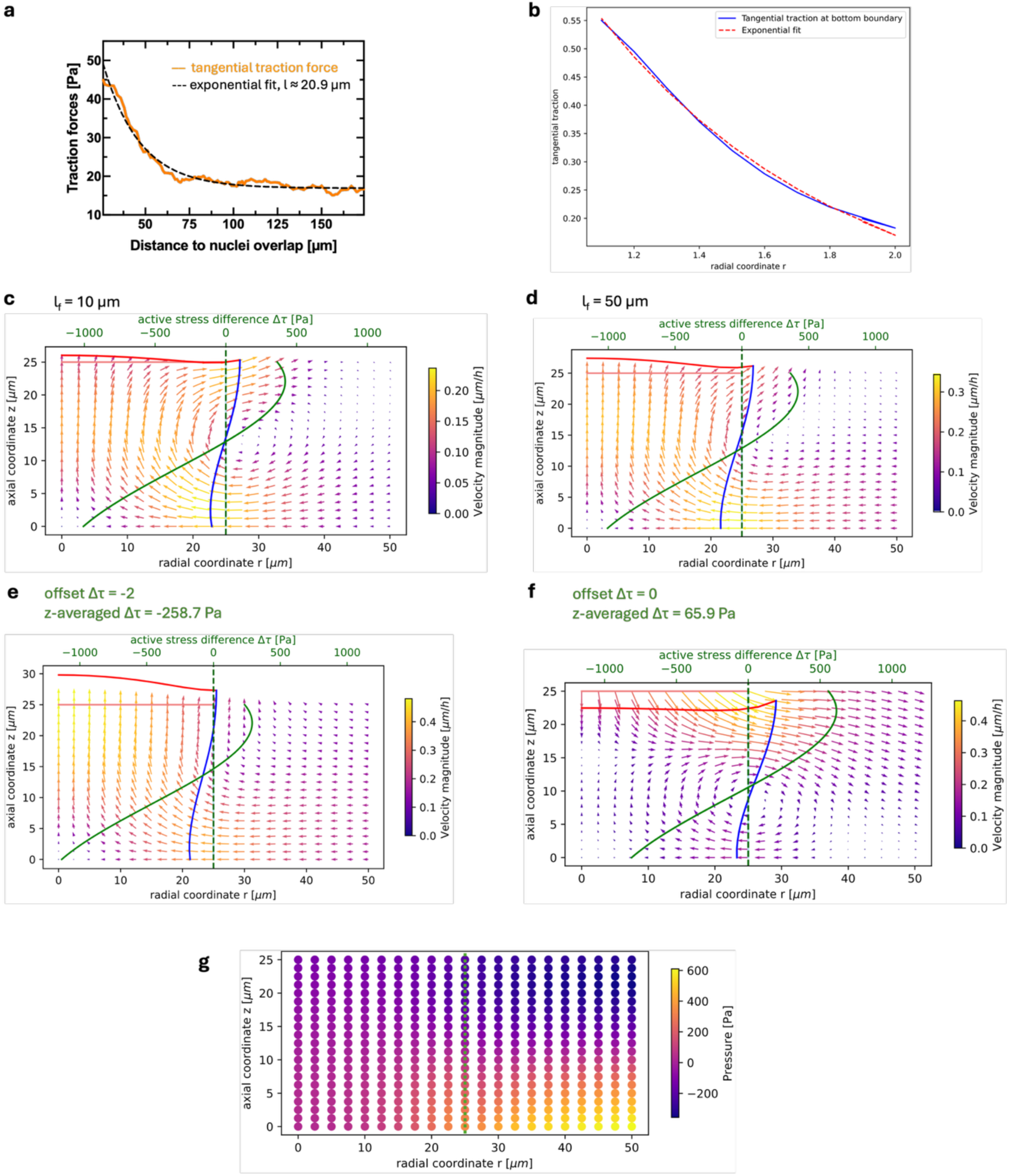
Numerical simulations of the model. **a** Experimentally measured basal tangential traction forces as a function of the distance to the center of the delaminating cell. Dotted lines show exponential fit with a characteristic length of 20.9 µm. **b** Tangential traction forces at the basal boundary in numerical simulations. **c** Simulated flow field obtained with friction length 10 µm. **d** Flow field with a friction length 50 µm. **e** Flow field with a increased (−2) or **f** reduced (0) offset of the active stress difference at the tissue boundary, changing the z-averaged active stress from negative to positive values. For **c** to **f**, note the changing flow fields and the differences in cell deformation, as highlighted by the blue boundary. **g** Z-dependence of the effective pressure in the delaminating cell and the surrounding tissue.

**Extended Data Figure 7:**
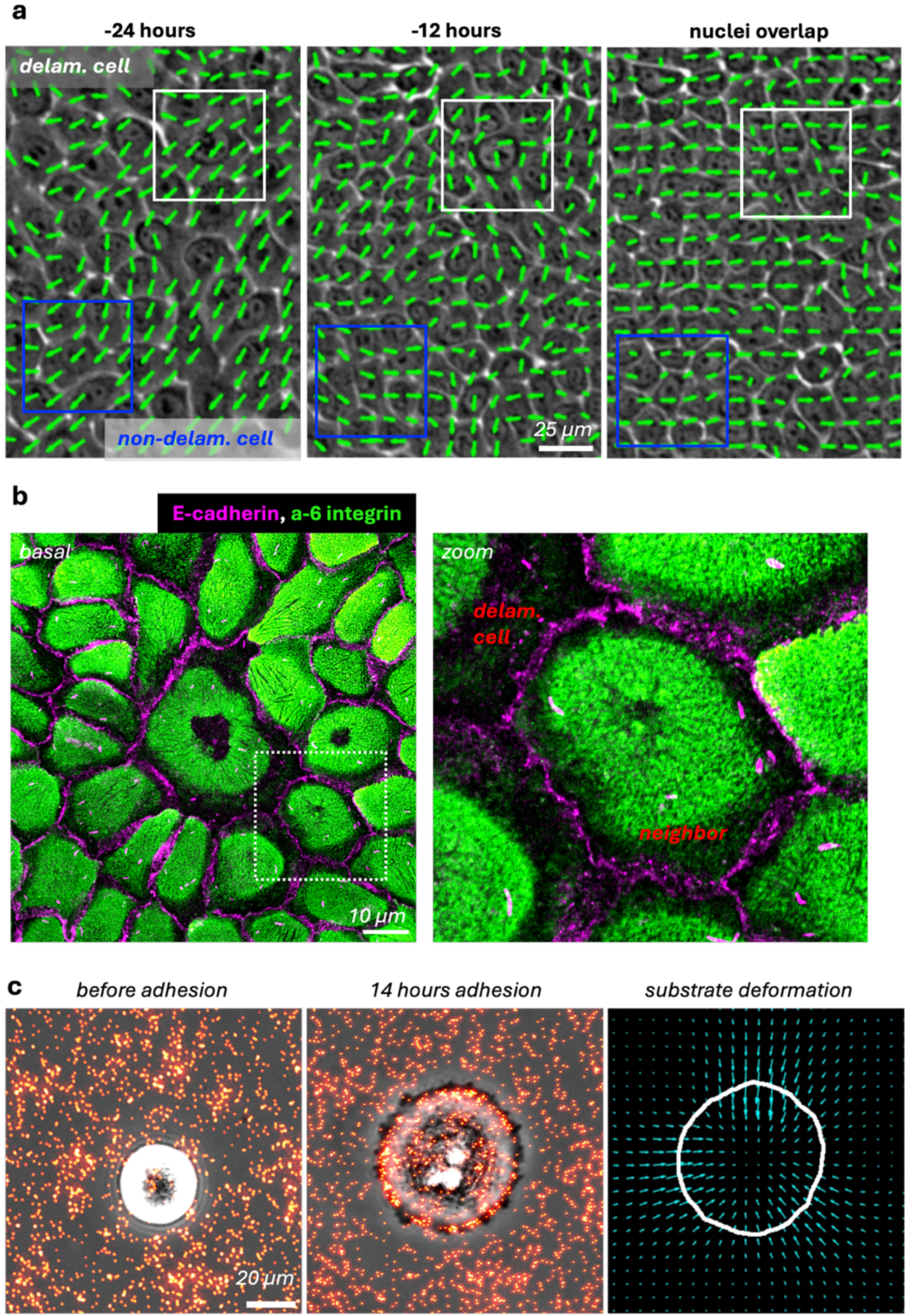
Convergent tissue flows. **a** Phase contrast images overlayed with the tissue flow orientation measured by Particle Image Velocimetry. Boxes show positions of a delaminating (white) and a non-delaminating (blue) cell, where the flow convergence was measured, related to Figure 6. **b** Confocal image of alpha-6 integrin (green) and E-cadherin (magenta) on the basal side. The central cell is in the process of delamination. Zoom-in highlight the absence of a polarity-and migration-indicating subcellular distribution of alpha-6 integrin. **c** Deformation of a 15kPa stiff E-cadherin-coated substrate containing fluorescent beads (red) by pre-differentiated cells. Left: bead distribution before cell adhesion. Middle: bead displacement 14 hours after adhesion. Right: Substrate deformation field. Cell outline in white.

**Extended Data Figure 8:**
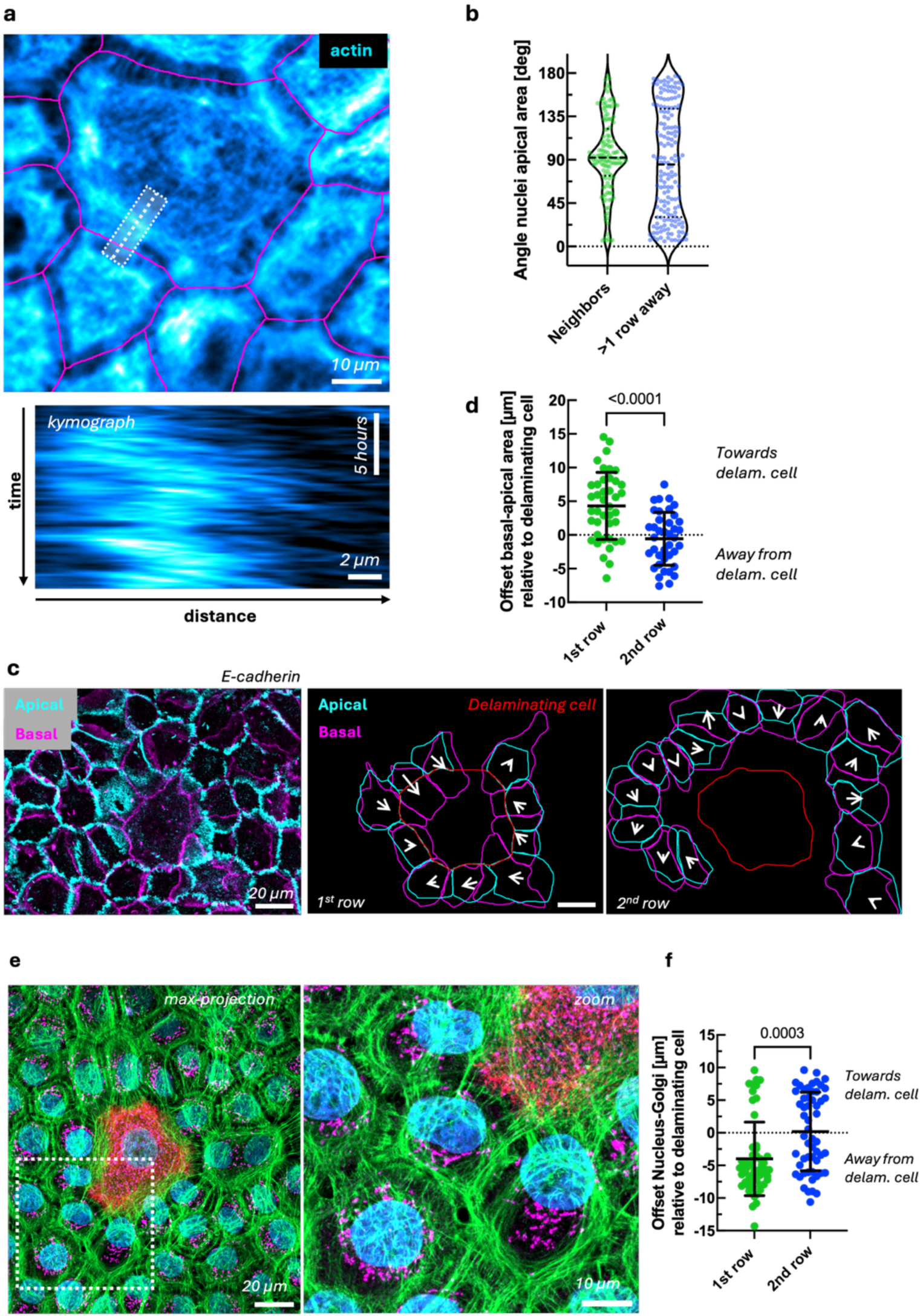
Actin retrograde flows pull on and reorient neighbor cells. **A** *Top*: Representative image of the actin cytoskeleton. Dotted line indicates position for the kymograph. *Bottom*: kymograph of the actin retrograde flow. **B** Distribution of the nuclei angles relative to the apical area of delaminating cells, related to Figure 6. Each dot represents 1 cell from N=3 measurements. **C** Images and **D** quantification of the offset between the basal and apical area of 1^st^ and 2^nd^ row neighbors. Apical (cyan) and basal (magenta) areas were segmented and the offset between the centers of mass was calculated. It was assigned a positive value, if the distance between the apical and the delaminating cell was greater than the distance between the basal and the delaminating cell. Each datapoint shows 1 cell. P-value from t-test. Means +/− SD. **E** Representative image of the Golgi apparatus (GM130: magenta, K10: red) at two days of differentiation. **F** Quantification of the offset between the nucleus and the Golgi relative to the position of differentiating and delaminating cells. Means +/− SD. P-value from t-test

