## Supplementary Information for "Local mechanical heterogeneity drives epidermal cell delamination"

**3 Numerical solution** **10**

**4 Discussion** **11**

**5 Generalization of the model to a nematic fluid** **12**

**A Some differential operator in cylindrical coordinates** **14**

### 1 Model

Motivated by the large time scales involved in the delamination process, we model the cell monolayer as an isotropic active low-Reynolds number fluid with viscosity $\eta$. We denote the fluid velocity as $\mathbf{v}(\mathbf{r})$ and the pressure as $p(\mathbf{r})$. We use standard cylindrical coordinates $(r, \theta, z)$ and unit basis $(\mathbf{e}_r, \mathbf{e}_\theta, \mathbf{e}_z)$, and assume axisymmetry around the $z$-axis, so that $\mathbf{v}$ and $p$ are independent of $\theta$, and $v^\theta = 0$. The problem is thus reduced to a two-dimensional domain in the $r$-$z$-plane with $r \in [0, L]$ and $z \in [0, h]$ where $L$ is the length of the domain in radial direction and $h$ the domain height.

The governing equations are

$$\nabla \cdot \mathbf{v} = 0 \tag{1}$$

$$\nabla \cdot \boldsymbol{\sigma} = \underline{0}. \tag{2}$$

In Eq. (1) we assume incompressibility and the force balance is given by Eq. (2). Herein, the total stress tensor is given by

$$\boldsymbol{\sigma} = 2\eta\mathbf{d} - p\mathbf{I} + \boldsymbol{\tau}, \tag{3}$$

where $\mathbf{d} = (\nabla\mathbf{v} + \nabla\mathbf{v}^\top)/2$ is the strain-rate tensor and $\mathbf{I}$ the identity matrix. $\boldsymbol{\tau}$ is the active stress. The delaminating cell is assumed to occupy the domain $r < r_c$ where $r_c(z, t)$ is the interface boundary between the delaminating cell and the surrounding tissue that depends on time $t$. We model the actomyosin contraction in the surrounding tissue by a constant prestress that is isotropic in each $(r,\theta)$-plane. With this the active stress reads in cylindrical coordinates

$$\boldsymbol{\tau}(r, z) = (\tau_0(z) + H(r - r_c)\Delta\tau(z)) \begin{bmatrix} 1 & 0 & 0 \\ 0 & 1 & 0 \\ 0 & 0 & 0 \end{bmatrix} . \tag{4}$$

Herein, $H(x)$ is the Heaviside step function, $\boldsymbol{\tau}_0(z)$ the stress field in the delaminating cell, and $\Delta\boldsymbol{\tau}(z)$ the active stress difference at the boundary between the delaminating cell and its neighbors. Note that $\boldsymbol{\tau}_0(z)$ only possesses horizontal components and only depends on $z$ so that its divergence vanishes everywhere. Hence only the jump part $\Delta\boldsymbol{\tau}(z)$ will matter in the dynamics and we set $\tau_0$ to zero for simplicity, keeping in mind that we describe the active stress up to a gauge $\boldsymbol{\tau}_0(z)$ and that the latter should be such that, according to experimental evidence, the total active stress $\boldsymbol{\tau}$ is everywhere positive. Below we will impose a $\Delta\boldsymbol{\tau}(z)$ that changes sign along the monolayer height, motivated by experimental observation.

We assume that the apical layer is free, implying vanishing normal stress at the top boundary of the domain. Motivated by the experimentally observed convergent tissue flows in the basal plane, we employ Navier-slip condition at the bottom boundary, allowing for partial sliding of the fluid with respect to the impermeable, underlying substrate. The boundary conditions thus are

$$(i) \quad v^z|_{z=0} = 0 \ ,$$

$$(ii) \quad \sigma^{zz}|_{z=h} = 0 \ ,$$

$$(iii) \quad \boldsymbol{\sigma}^{||z}|_{z=0} = -\alpha \mathbf{v}^{||} \ ,$$

$$(iv) \quad \boldsymbol{\sigma}^{||z}|_{z=h} = 0 \ ,$$

where $||$ denotes the projection on the horizontal $(r,\theta)$-plane and $\alpha = \eta/l_f$ is the tangential friction coefficient with the friction length $l_f$. We solve Eqs. (1) and (2) for the velocity $\mathbf{v} = v^r(r,z)\mathbf{e}_r + v^z(r,z)\mathbf{e}_z$ and the pressure $p(r,z)$ analytically in section 2 and numerically in section 3.

The problem can be made dimensionless by choosing as characteristic scales $h$ for length, $\eta/\Delta\tau_m$ for time, and $\Delta\tau_m$ for pressure, where $\Delta\tau_m = \max|\Delta\tau(z)|$. We find that the only remaining non-dimensional parameter is $\Lambda = h\alpha/\eta = h/l_f$ that compares the monolayer height with the slip length. This means that the solution for the velocity field depends only on $\Lambda$, i.e., one has $|\mathbf{v}| = \frac{h\Delta\tau_m}{\eta} g(\Lambda)$ where $g(\Lambda)$ is a dimensionless function describing how friction with the substrate modifies the velocity magnitude.

### 2 Analytical approach

In this section, we consider a slightly modified problem that we solve analytically. First, we assume that the delaminating cell has initially a cylindrical shape of radius $r_c = R$ and we will solve only for the initial velocity field of the delamination process. We further assume the system to be infinite in the radial direction, *i.e.* $L \to \infty$, and that the velocity remains bounded at $r = 0$ and $r \to \infty$. This constitute the fifth boundary condition of our analytical approach:

$$(v) \quad \lim_{r \to 0,\infty} |\mathbf{v}| < \infty \ .$$

Moreover, we will solve separately the dynamics inside and outside the delaminating cell, and assume continuity of the stress at the interface, *i.e.*

$$(vi) \quad \sigma \cdot \mathbf{e}_r|_{r=R^-} = \sigma \cdot \mathbf{e}_r|_{r=R^+} \ .$$

#### 2.1 Exploiting the rotational symmetry

The unknown variables of the model described in the previous section are the velocity and pressure fields: $(\mathbf{v}, p)$. In this section, we leverage the cylindrical symmetry of our problem to reformulate it in terms of a single scalar function: the so-called Stokes stream function.

##### 2.1.1 The Stokes stream function and its dynamics

Since we assume incompressibility, there exists a vector field $\mathbf{A}$ such that

$$\mathbf{v} = \nabla \times \mathbf{A} \ . \tag{5}$$

The rotational invariance further implies that we can assume $\mathbf{A}$ to be purely along $\mathbf{e}_\theta$. Indeed, if $\mathbf{A}(r,z) = A^r\mathbf{e}_r + A^\theta\mathbf{e}_\theta + A^z\mathbf{e}_z$, then (see appendix A for useful formulas in cylindrical coordinates):

$$\nabla \times \mathbf{A} = -\partial_z A^\theta \mathbf{e}_r + \underbrace{(\partial_z A^r - \partial_r A^z)}_{=0}\mathbf{e}_\theta + \frac{1}{r}\partial_r(rA^\theta)\mathbf{e}_z \ . \tag{6}$$

Hence we can make the gauge choice $A^r = A^z = 0$ and define the so-called "Stokes stream function", $\psi(r,z) \equiv rA^\theta$, so that

$$v^r = -\tfrac{1}{r}\partial_z\psi \tag{7}$$

$$v^z = \tfrac{1}{r}\partial_r\psi \tag{8}$$

In turn, taking the curl of the Stokes equation is readily shown to be equivalent to the PDE

$$\mathcal{L}^2\psi = 0 \tag{9}$$

with the operator

$$\mathcal{L} \equiv \partial_z^2 + \partial_r^2 - \frac{1}{r}\partial_r \ . \tag{10}$$

##### 2.1.2 Pressure, stress, and boundary conditions in terms of $\psi$

Since the velocity field is divergence free, $\Delta\mathbf{v} = -(\nabla\times)^2\mathbf{v}$. Then, using (5) and (60), we get

$$\Delta\mathbf{v} = \nabla \times \left(\frac{1}{r}\begin{bmatrix} 0 \\ \mathcal{L}\psi \\ 0 \end{bmatrix}\right) \ . \tag{11}$$

Finally, using eq. (58), with $\psi$ replaced by $\mathcal{L}\psi$, we conclude that the Stokes equation (2) can be re-written in terms of $\psi$ as:

$$\partial_r p = -\frac{\eta}{r}\partial_z\mathcal{L}\psi \ , \tag{12}$$

$$\partial_z p = \frac{\eta}{r}\partial_r\mathcal{L}\psi \ . \tag{13}$$

As for the stress tensor, it reads

$$\sigma = \begin{bmatrix} -2\eta\partial_r\frac{1}{r}\partial_z\psi - p + \tau & 0 & \eta(\partial_r\frac{1}{r}\partial_r\psi - \frac{1}{r}\partial_z^2\psi) \\ 0 & \frac{-2\eta}{r^2}\partial_z\psi - p + \tau & 0 \\ \eta(\partial_r\frac{1}{r}\partial_r\psi - \frac{1}{r}\partial_z^2\psi) & 0 & 2\eta\frac{1}{r}\partial_r\partial_z\psi - p \end{bmatrix} \ . \tag{14}$$

Finally, the boundary conditions can also be reformulated using the stream function[1]

$$(i) \quad \partial_r\psi|_{z=0} = 0 \ ,$$

[1]Note that conditions *(i)* and *(ii)*, that read $\partial_r\psi(r, z = 0, h) = 0$, are satisfied for all $r$. We can thus take the derivative of these equalities with respect to $r$, leading to $\partial_r^2\psi(r, z = 0, h) = 0$.

$$(ii) \quad \left[\frac{2\eta}{r}\partial_r\partial_z\psi - p\right]\Big|_{z=h} = 0 \ ,$$

$$(iii) \quad \left[\partial_z^2\psi + \frac{\alpha}{\eta}\partial_z\psi\right]\Big|_{z=0} = 0 \ ,$$

$$(iv) \quad \partial_z^2\psi|_{z=h} = 0 \ ,$$

$$(v) \quad \lim_{r\to 0,\infty} \frac{1}{r^2}[(\partial_r\psi)^2 + (\partial_z\psi)^2] < \infty \ ,$$

$$(vi) \quad \begin{bmatrix} -2\eta\partial_r\frac{1}{r}\partial_z\psi - p + \tau \\ 0 \\ \eta(\partial_r\frac{1}{r}\partial_r\psi - \frac{1}{r}\partial_z^2\psi) \end{bmatrix}\Bigg|_{r=R^-} = \begin{bmatrix} -2\eta\partial_r\frac{1}{r}\partial_z\psi - p + \tau \\ 0 \\ \eta(\partial_r\frac{1}{r}\partial_r\psi - \frac{1}{r}\partial_z^2\psi) \end{bmatrix}\Bigg|_{r=R^+} .$$

#### 2.2 A basis of solutions separated in $r$ and $z$ to the free-boundary problem

We now look for solutions of the free-boundary problem associated to eq. (9) which are of the form:

$$\psi_k(r,z) = \chi_k(r)\phi_k(z) \ , \tag{15}$$

with $k \in \mathbb{C}$. We assume that $\chi_k$ is in the kernel of the operator

$$\mathcal{L}_{r,k} \equiv \partial_r^2 - \frac{1}{r}\partial_r - k^2 \ , \tag{16}$$

and look for its corresponding explicit expression.

- For the case $k \neq 0$, we make the change of variable $\chi_k(r) \to y_k(rk) \equiv \chi_k(r)/r$. The equation $\mathcal{L}_{r,k}\chi_k = 0$ becomes $x^2y''(x) + xy'(x) - (1 + x^2)y(x) = 0$, with $x = rk$, which is a modified Bessel equation whose two-dimensional solution space is spanned by the modified Bessel functions of first and second kind and first order: $I_1(x)$ and $K_1(x)$. Going back to the initial variable and function, we thus get:

$$\ker(\mathcal{L}_{r,k}) = \text{Span}\left\{rI_1(kr), rK_1(kr)\right\} \ , \tag{17}$$

- For the $k = 0$ case, the space of solutions is readily shown to read:

$$\ker(\mathcal{L}_{r,k=0}) = \text{Span}\left\{1, r^2\right\} \ . \tag{18}$$

We now turn to the identification of $\phi_k(z)$. Because $\chi_k \in \ker\mathcal{L}_{r,k}$ and $\psi_k = \chi_k\phi_k$ satisfies equation (9), then $\phi_k \in \ker(\mathcal{L}_{k,z}^2)$, where $\mathcal{L}_{k,z} \equiv \partial_z^2 + k^2$. Again, we need to distinguish the same two cases:

- $k \neq 0$: we know $\ker(\mathcal{L}^2_{k,z})$ is 4-dimensional, and that it contains $\ker \mathcal{L}_{k,z} = \text{Span}\{\cos(kz), \sin(kz)\}$. It further can be shown, by a direct calculation, that $z \mapsto z\cos(kz)$ and $z \mapsto z\sin(kz)$ span the two missing dimensions. Thus

$$\ker\left(\mathcal{L}^2_{k,z}\right) = \text{Span}\{\cos(kz), \sin(kz), z\cos(kz), z\sin(kz)\} \ . \tag{19}$$

- $k = 0$: the solution is now obviously

$$\ker\left(\mathcal{L}^2_{k=0,z}\right) = \ker \partial_z^4 = \text{Span}\left\{1, z, z^2, z^3\right\} \ . \tag{20}$$

#### 2.3 A specified basis for apical and basal boundary conditions

In this section, we specify those functions of the form (15) that, in addition of solving eq. (9), satisfy boundary conditions *(i)*–*(iv)*. To this aim, we start by reformulating condition *(iv)* in terms of $\psi$ only. Note that this condition, which reads $(2\eta/r)\partial_r\partial_z\psi - p(r, z = h) = 0$ in terms of $\psi$ *and* the pressure $p$, must be valid for all $r$, and can thus be derived with respect to $r$. Doing so and using (12), we get a condition that we denote $\partial_r(ii)$ and that reads

$$\left[2\eta\partial_z\partial_r\frac{1}{r}\partial_r\psi + \frac{\eta}{r}\partial_z\mathcal{L}\psi\right]\Bigg|_{z=h} = 0 \ , \tag{21}$$

*i.e.*

$$\partial_z\left[3(\partial_r^2\psi - \frac{1}{r}\partial_r\psi) + \partial_z^2\psi\right]\Bigg|_{z=h} = 0 \ . \tag{22}$$

We now turn to determining the basis of solutions to eq. (9) that is adapted to conditions *(i)*, $\partial_r(ii)$, *(iii)*, and *(iv)*, and we will see later what additional condition *(ii)* imposes on the global solution compared to $\partial_r(ii)$.

##### 2.3.1 Basis functions for $k \neq 0$

We look for a solution $\psi_k(r, z)$ to eq. (9) of the form (15), with $\chi_k \in \ker(\mathcal{L}_{r,k})$ and $\phi_k \in \ker(\mathcal{L}^2_{k,z})$ that reads

$$\phi_k(z) = (A_k + B_k z)\sin(kz) + (C_k + D_k z)\cos(kz) \ . \tag{23}$$

Injecting $\psi_k$ into conditions *(i)*, $\partial_r(ii)$, *(iii)*, and *(iv)* allows expressing these conditions in terms of $\phi_k$ only:

$$(i) \qquad \phi_k(0) = 0 \ , \tag{24}$$

$$\partial_r(ii) \qquad 3k^2\phi_k'(h) + \phi_k^{(3)}(h) = 0 \ , \tag{25}$$

$$(iii) \qquad \phi_k''(0) + \frac{\alpha}{\eta}\phi_k'(0) = 0 \ , \tag{26}$$

$$(iv) \qquad \phi_k''(h) - k^2\phi_k(h) = 0 \ . \tag{27}$$

The first condition directly gives $C_k = 0$. Straightforward calculations then allow translating the three other conditions in terms of the constants $A_k, B_k$, and $D_k$ as:

$$\begin{bmatrix} \cos(kh) & h\cos(kh) & -h\sin(kh) \\ k & 2\eta k/\alpha & 1 \\ k\sin(kh) & kh\sin(kh)-\cos(kh) & kh\cos(kh)+\sin(kh) \end{bmatrix} \begin{bmatrix} A_k \\ B_k \\ D_k \end{bmatrix} = 0 \ . \quad (28)$$

In order for a solution to exist, the determinant of the matrix on the left-hand side of this last equation must vanish. This reads

$$\frac{2\eta k}{\alpha}[kh + \cos(kh)\sin(kh)] + \cos^2(kh) - k^2h^2 = 0 \quad (29)$$

For $k$ satisfying eq. (29), the space of solutions can be shown to be generated by the vector

$$\begin{bmatrix} A_k \\ B_k \\ D_k \end{bmatrix} = \begin{bmatrix} kh^2 \\ -kh - \sin(kh)\cos(kh) \\ -\cos^2(kh) \end{bmatrix} \ . \quad (30)$$

Note that, the dominating term of the left-hand side of eq. (29) for large $k$ is $k^2h^2(2\eta/\alpha h - 1)$. Hence, unless $\eta/h\alpha = 1/2$, the left hand-side of eq. (29) is strictly positive for real, sufficiently large $k$'s. This mean that we generically need to consider complex $k$'s in order for $(\psi_k)_k$ to be a basis of solutions. We denote by $\mathcal{S}$ the (complex) set of solutions of eq. (29) and call it the spectrum of the problem. Because eq. (29) is invariant under both complex conjugation and $k \mapsto -k$, the spectrum $\mathcal{S}$ is symmetric with respect to both imaginary and real axis in the complex plane.

###### 2.3.2 Basis functions for $k = 0$

For $k = 0$, as we have seen in section 2.2, the general solution of eq. (9) reads:

$$\psi_0(r,z) = A_1 + B_1 z + \frac{C_1}{2}z^2 + \frac{D_1}{3}z^3 + r^2(A_2 + B_2 z + \frac{C_2}{2}z^2 + \frac{D_2}{3}z^3) \ , \quad (31)$$

with $A_i, B_i, C_i, D_i, i \in \{1,2\}$, real constants. In this case, conditions *(i)*, $\partial_r$*(ii)*, *(iii)*, and *(iv)* read

$$(i) \quad A_2 = 0 \ , \quad (32)$$

$$\partial_r(ii) \quad D_1 + D_2 r^2 = 0 \ \forall r \Rightarrow D_1 = D_2 = 0 \quad , \quad (33)$$

$$(iii) \quad C_1 + \frac{\alpha}{\eta}B_1 + (C_2 + \frac{\alpha}{\eta}B_2)r^2 = 0 \ \forall r \Rightarrow C_1 + \frac{\alpha}{\eta}B_1 = C_2 + \frac{\alpha}{\eta}B_2 = 0 \ , \quad (34)$$

$$(iv) \quad C_1 + 2D_1 h + (C_2 + 2D_2 h)r^2 = 0 \ \forall r \Rightarrow C_1 + 2D_1 h = C_2 + 2D_2 h = 0 \ . \quad (35)$$

These constraints imply that all the $A_i, B_i, C_i, D_i$ vanish, except $A_1$, and consequently $\psi_{k=0}$ is a constant.

##### 2.3.3 General solution

The general solution to eq. (9), that satisfies boundary conditions *(i)*, $\partial_r(ii)$, *(iii)*, and *(iv)*, thus reads

$$\psi(r,k) = c_0 + \sum_{k\in\mathcal{S}} c_k \chi_k(r)\phi_k(z) \ , \tag{36}$$

where all $c_k$'s are complex constants and $\phi_k$ is given by eqs. (23) & (30). As for $\chi_k(r)$, taking into account the boundedness condition *(v)* implies that it is respectively equal to $rI_1(|k|r)$ and $rK_1(|k|r)$ in the delaminating cell $r < R$ and in the rest of the tissue, where $r > R$.

Then, injecting (36) into eqs. (12)-(13) and integrating the resulting vector field leads to the following expression for the pressure:

$$p(r,z) = p_0 + \frac{2\eta}{r} \sum_{k\in\mathcal{S}} c_k \chi_k'(r) \left[B_k \sin(kz) + D_k \cos(kz)\right] \ , \tag{37}$$

where $p_0$ is a constant. Note that injecting (36) & (37) into condition *(ii)* gives $p_0 = 0$, which is the part of the condition we missed by considering $\partial_r(ii)$ instead of the full *(ii)* earlier.

#### 2.4 Incorporating the last ingredient: the active stress jump

We finally need to take into account condition *(vi)*, that imposes action -reaction between the delaminating cell and the rest of the tissue. Denoting respectively with exponents D and T the quantities that relate to the delaminating cell and the tissue, we first get that the continuity of the tangential stress gives

$$c_k^D \chi_k^D(R) = c_k^T \chi_k^T(R) \ . \tag{38}$$

Taking this into account, the continuity of the normal stress then gives

$$\Delta\tau(z) = -\frac{2\eta}{R} \sum_{k\in\mathcal{S}} c_k^D \left[\chi_D' - \frac{\chi_D}{\chi_T}\chi_T'\right]\Bigg|_{r=R} f_k(z) \ , \tag{39}$$

where $\Delta\tau \equiv \tau^T - \tau^D$, $\left[\chi_D' - \chi_D\chi_T'/\chi_T\right](R) = Rk\left[I_0 + K_0 I_1/K_1\right](kR) > 0$, and $f_k(z) \equiv \left[(kA_k + 2D_k + kB_k z)\cos(kz) + (2B_k - kD_k z)\sin(kz)\right]$.

Thus, for a given jump of active stress $\Delta\tau(z)$, we obtain the coefficients $c_k^D$, thanks eq. (39), by projecting $\Delta\tau$ over the basis functions $f_k$. Then, eq. (38) straightforwardly gives the coefficients $c_k^T$. We can consequently deduce the corresponding Stokes stream functions $\psi$ given by eq. (36) and, in turn, the velocity and pressure fields inside and outside the delaminating cell, thanks to eqs. (7), (8), (12), & (13).

We are now in position to (qualitatively) demonstrate that a distribution of active stress that is dominated respectively by the delaminating cell on the basal side and the rest of the tissue on the apical side does initiate the delamination of the central cell. To this end, we could choose any arbitrary profile $\Delta\tau(z)$ that is negative for small $z$ and becomes positive as $z$ increases up to $h$. In order to avoid explicitly performing the projection of $\Delta\tau(z)$ onto the "eigen-stress" basis $f_k$, we directly pick an appropriate eigen-stress $f_{k_0}$ and choose its amplitude as to match the typical speed observed experimentally, $v \sim 0.5\mu m.h^{-1}$. For the physical parameters measured on the experimental system (see main text for details), we choose the imaginary part of $f_{k_0}$ with $k_0 \in \mathcal{S}$ the eigenvalue in the positive quadrant of $\mathbb{C}$ with the second smallest real part. We see that such (eigen-) active stress displays the overall shape we want and indeed generates a flow that is physically coherent with the earlier phase of the delamination process (see Figure 5b of the main text).

### 3 Numerical solution

In our numerical approach we assume that the problem domain is along the radial direction embedded in adjacent tissue that is subject to the same active prestress. We therefore write as boundary condition

$$\boldsymbol{\sigma}\mathbf{e}_r|_{r=L} = \boldsymbol{\tau}\mathbf{e}_r|_{r=L} \tag{40}$$

with $\boldsymbol{\tau}$ given by Eq. (4). We use a single simulation domain and therefore the velocity is continuous at the delaminating cell boundary.

Assuming a stationary vertical interface, we have $r_c(z,t) = R$ and $\boldsymbol{n} = \boldsymbol{e}_r$. Choosing $\tau_0(z) = 0$, the active stress in Eq. (4) is implemented as

$$\boldsymbol{\tau} = H_w(r - r_c)\Delta\tau(z) \begin{bmatrix} 1 & 0 & 0 \\ 0 & 1 & 0 \\ 0 & 0 & 0 \end{bmatrix}. \tag{41}$$

Herein, we use a smoothed step function

$$H_w(x) = \frac{1}{2}\left[1 + \tanh\left(\frac{x}{w}\right)\right] \tag{42}$$

where $w$ is the width of the interface between the delaminating cell and its neighbors. The numerical approach allows us to impose an arbitrary function for $\Delta\tau(z)$. We decompose the stress jump in Eq. (41) according to

$$\Delta\tau(z) = \langle\Delta\tau\rangle_z + \delta\Delta\tau(z), \tag{43}$$

where $\langle\Delta\tau\rangle_z$ is the constant part with $\langle.\rangle_z$ being the $z$-average over the height of the domain and $\delta\Delta\tau(z)$ denotes the apico-basal variation of active stress difference with $\langle\delta\Delta\tau(z)\rangle_z = 0$. From Eq. (43) we define the ratio $\varepsilon_\tau = \frac{\max|\delta\Delta\tau(z)|}{\langle\Delta\tau(z)\rangle_z}$ which represents a second dimensionless parameter in the problem in addition to the length ratio $\Lambda$. We choose $\delta\Delta\tau(z)$ fixed and equal to the deviation of the average stress difference in our analytical approach, see Eq. (39). We vary $\varepsilon_\tau$ by changing $\langle \Delta\tau \rangle_z$ in our simulations to model the flow field for different prestress differences between the delaminating cell and the neighboring tissue.

We solve Eqs. (1) and (2) together with the boundary conditions detailed above using the finite element method. To this end we compute the weak form of the governing equations which is then implemented in `fenicsx`. The computational domain is partitioned into triangular elements where $N$ is the number of elements and $h_t$ their characteristic length. We employ Taylor–Hood finite elements, using cubic piecewise polynomial basis functions for the velocity and quadratic basis functions for the pressure. We choose the interface width so that $h_t/2 < w \ll R$ to ensure appropriate resolution of the interface on the mesh. We use the parameters listed in table 1.

| Parameter: | Value: |
|---|---|
| fluid viscosity $\eta$ | 25 |
| interface position $R$ | 25 |
| outer boundary position $L$ | 75 |
| domain height $h$ | 25 |
| Navier friction coefficient $\alpha$ | 1 |
| number of mesh vertices $N$ | 441 |
| interface width $w$ | 2.5 |

Table 1: Simulation parameters.

Our numerical simulations give access to the traction force at the basal layer which decays with the distance from the delaminating cell (ED Fig. 6b). This is in alignment with experiments (ED Fig. 6a). Comparison of the experimental curve yields a characteristic pressure scale. Assuming a delaminating cell size of $R = 25\,\mu\text{m}$ and a velocity of $0.5\,\mu\text{m}\,\text{h}^{-1}$ in alignment with typical experimental values, this yields a fluid viscosity of $\eta_{\text{eff}} \approx 4 \times 10^3\,\text{Pa}\,\text{h}$.

### 4 Discussion

In our numerical approach the tangential velocity is continuous at the delaminating cell boundary, modelling a classical fluid-fluid interface. In our analytical approach, to help solving the problem explicitly, we allow for a discontinuity of the tangential velocity across the interface, which introduces tangential slip at both sides of the boundary. In both approaches the normal component of the velocity is continuous across the interface, consistent with the kinematic condition for an impermeable boundary. Furthermore, our analytical and numerical approaches differ in the applied boundary conditions on the outer radial domain boundary. In the analytical approach we impose that the velocity remains finite, while in our numerical simulations we impose a more specific continuity of the active stress. Despite these distinct modelling choices, both approaches produce similar trends for velocity field, leading to cellular upward motion and shape deformation that is in alignment with experimental observations.

### 5 Generalization of the model to a nematic fluid

In the following we discuss the generalization of our analysis to the case of a nematic fluid. The active stress in the tissue surrounding the delaminating cell (for $r > r_c$) is equivalent to the superposition of an isotropic tension $\frac{2\Delta\tau(z)}{3}\mathbf{I}$ and a traceless tensor given by

$$\mathbf{Q} = \begin{bmatrix} 1 & 0 & 0 \\ 0 & 1 & 0 \\ 0 & 0 & -2 \end{bmatrix} . \tag{44}$$

Eq. (44) describes a nematic order parameter $Q_{\alpha\beta} = \langle n_\alpha n_\beta - \frac{1}{3}\delta_{\alpha\beta}\rangle$ that corresponds to filaments that are aligned parallel to the $z$-axis and disordered in the $r$-$\theta$-plane. We can therefore write the stress as

$$\boldsymbol{\sigma} = -p'\mathbf{I} + 2\eta\mathbf{d} + \frac{\Delta\tau(z)}{3}\mathbf{Q} \tag{45}$$

where

$$p' = p - \frac{2\Delta\tau(z)}{3} \tag{46}$$

is the modified pressure. The standard evolution equation for the nematic order parameter is given by

$$\frac{D\mathbf{Q}}{Dt} = \beta\mathbf{d} - \Gamma\frac{\delta\mathcal{F}}{\delta\mathbf{Q}} \tag{47}$$

where $\beta$ is the flow alignment coefficient, $\Gamma$ the mobility, and

$$\mathcal{F} = \frac{1}{2}\int \left[\mathbf{Q}^2 + \xi^2\left(\nabla\mathbf{Q}\right)^2\right] d^3r \tag{48}$$

is the free energy. We assume that the nematic order parameter is dynamically slaved to the velocity and therefore the time-derivative in Eq. (47) vanishes. We can therefore write using Eqs. (48) and (47)

$$\beta\mathbf{d} = \Gamma\left(\mathbf{Q} + \xi^2\nabla^2\mathbf{Q}\right) . \tag{49}$$

Neglecting spatial gradients in the nematic tensor, we obtain

$$\mathbf{Q} = \frac{\beta}{\Gamma}\mathbf{d} \tag{50}$$

and Eq. (45) becomes

$$\boldsymbol{\sigma} = p'\mathbf{I} + 2\eta'\mathbf{d} \tag{51}$$

where

$$\eta' = \eta\left(1 + \frac{\beta}{6\Gamma}\Delta\tau(z)\right) \tag{52}$$

is the modified viscosity. This means that the active stress can be fully absorbed into a passive stress with a renormalized pressure and viscosity.

From Eq. (50) follows that $Q_{r\theta} = Q_{\theta r} = Q_{\theta z} = Q_{z\theta}$ and thus the azimuthal direction is a principal axis of the nematic tensor. The eigenvalue along this axis is given by $\lambda_\theta = \frac{\beta v_r}{\Gamma r}$. The two other principal axes lie in the $r$-$z$-plane. The eigenvalues along these axes are the roots of the characteristic polynomial

$$\det\left[\frac{\beta}{\Gamma}\begin{pmatrix} d_{rr} & d_{rz} \\ d_{zr} & d_{zz} \end{pmatrix} - \lambda \mathbf{I}\right] = 0 \tag{53}$$

where $d_{rr} = \partial_r v^r$, $d_{rz} = d_{zr} = \frac{1}{2}(\partial_r v^z + \partial_z v^r)$, and $d_{zz} = \partial_z v^z$ are the entries of the strain-rate tensor. We obtain

$$\lambda_\pm = \frac{\beta}{2\Gamma}\left[(d_{rr} + d_{zz}) \pm \sqrt{(d_{rr} - d_{zz})^2 + 4d_{rz}^2}\right]. \tag{54}$$

A ring-like nematic order occurs when the azimuthal eigenvalue is the largest eigenvalue of $\mathbf{Q}$. This is the case for

$$\frac{v_r}{r} > \frac{1}{2}\left[(d_{rr} + d_{zz}) + \sqrt{(d_{rr} - d_{zz})^2 + 4d_{rz}^2}\right]. \tag{55}$$

### A Some differential operator in cylindrical coordinates

The gradient of a vector in cylindrical coordinates reads

$$\nabla \mathbf{v} = \begin{bmatrix} \partial_r v^r & \frac{1}{r}(\partial_\theta v^r - v^\theta) & \partial_z v^r \\ \partial_r v^\theta & \frac{1}{r}(\partial_\theta v^\theta + v^r) & \partial_z v^\theta \\ \partial_r v^z & \frac{1}{r}\partial_\theta v^z & \partial_z v^z \end{bmatrix} . \tag{56}$$

Assuming rotation invariance, the strain-rate tensor is then

$$\mathbf{d} = \frac{1}{2} \begin{bmatrix} 2\partial_r v^r & 0 & \partial_r v^z + \partial_z v^r \\ 0 & 2v^r/r & 0 \\ \partial_r v^z + \partial_z v^r & 0 & 2\partial_z v^z \end{bmatrix} . \tag{57}$$

Successive actions of the curl operator:

- By definition of the stream function and the corresponding vector field $\mathbf{A}$:

$$\nabla \times \mathbf{A} = \nabla \times \begin{bmatrix} 0 \\ \psi/r \\ 0 \end{bmatrix} = \frac{1}{r} \begin{bmatrix} -\partial_z \psi \\ 0 \\ \partial_r \psi \end{bmatrix} \tag{58}$$

- using the formula of the curl operator in the (unit) cylindrical coordinate system, which reads for a generic vector field $\mathbf{U}$

$$\nabla \times \mathbf{U} = \begin{bmatrix} \frac{1}{r}\partial_\theta U^z - \partial_z U^\theta \\ \partial_z U^r - \partial_r U^z \\ \frac{1}{r}\partial_r (r U^\theta) - \frac{1}{r}\partial_\theta U^r \end{bmatrix} \tag{59}$$

  a straightforward calculation gives

$$(\nabla \times)^2 \mathbf{A} = (\nabla \times)^2 \begin{bmatrix} 0 \\ \psi/r \\ 0 \end{bmatrix} = -\frac{1}{r} \begin{bmatrix} 0 \\ \mathcal{L}\psi \\ 0 \end{bmatrix} . \tag{60}$$

Using the definition of $\mathbf{A}$ and $\psi$, together with eq. (60) – twice applied – and the fact that, for the divergence-free vector field $\mathbf{v}$, $\Delta \mathbf{v} = -(\nabla \times)^2 \mathbf{v}$, we get

$$\nabla \times \Delta \mathbf{v} = -\frac{1}{r} \begin{bmatrix} 0 \\ \mathcal{L}^2 \psi \\ 0 \end{bmatrix} . \tag{61}$$
